# Structural classification of neutralizing antibodies against the SARS-CoV-2 spike receptor-binding domain suggests vaccine and therapeutic strategies

**DOI:** 10.1101/2020.08.30.273920

**Authors:** Christopher O. Barnes, Claudia A. Jette, Morgan E. Abernathy, Kim-Marie A. Dam, Shannon R. Esswein, Harry B. Gristick, Andrey G. Malyutin, Naima G. Sharaf, Kathryn E. Huey-Tubman, Yu E. Lee, Davide F. Robbiani, Michel C. Nussenzweig, Anthony P. West, Pamela J. Bjorkman

## Abstract

The COVID-19 pandemic presents an urgent health crisis. Human neutralizing antibodies (hNAbs) that target the host ACE2 receptor-binding domain (RBD) of the SARS-CoV-2 spike^1–5^ show therapeutic promise and are being evaluated clincally^6–8^. To determine structural correlates of SARS-CoV-2 neutralization, we solved 8 new structures of distinct COVID-19 hNAbs^5^ in complex with SARS-CoV-2 spike trimer or RBD. Structural comparisons allowed classification into categories: (1) *VH3-53* hNAbs with short CDRH3s that block ACE2 and bind only to “up” RBDs, (2) ACE2-blocking hNAbs that bind both “up” and “down” RBDs and can contact adjacent RBDs, (3) hNAbs that bind outside the ACE2 site and recognize “up” and “down” RBDs, and (4) Previously-described antibodies that do not block ACE2 and bind only “up” RBDs^9^. Class 2 comprised four hNAbs whose epitopes bridged RBDs, including a *VH3-53* hNAb that used a long CDRH3 with a hydrophobic tip to bridge between adjacent “down” RBDs, thereby locking spike into a closed conformation. Epitope/paratope mapping revealed few interactions with host-derived *N*-glycans and minor contributions of antibody somatic hypermutations to epitope contacts. Affinity measurements and mapping of naturally-occurring and in vitro-selected spike mutants in 3D provided insight into the potential for SARS-CoV-2 escape from antibodies elicited during infection or delivered therapeutically. These classifications and structural analyses provide rules for assigning current and future human RBD-targeting antibodies into classes, evaluating avidity effects, suggesting combinations for clinical use, and providing insight into immune responses against SARS-CoV-2.

---

Neutralizing antibodies (NAbs) against SARS-CoV-2 protect against infection in animal models^1,3,4,10,11^ and are being evaluated for prophylaxis and as therapeutics in humans^7,8^. These antibodies target the SARS-CoV-2 spike (S) trimer^3,5,10,12–17^, a viral glycoprotein that mediates binding to angiotensin-converting enzyme 2 (ACE2) receptor^18,19^. S trimer comprises three copies of an S1 subunit containing the receptor-binding domain (RBD) and three copies of S2, which includes the fusion peptide and transmembrane regions^20,21^. The RBDs of SARS-CoV-2 and other coronaviruses exhibit flexibility, such that they bind ACE2 only when they are in an “up” conformation, as compared with the “down” RBD conformation of the closed, prefusion S trimer^20–25^.

Many hNAbs isolated from COVID-19 convalescent donors target the RBD, binding to distinct, sometimes non-overlapping, epitopes^3–5,10,12–14,17^. A subset of these antibodies blocks viral entry by binding to the ACE2-binding site on the RBD^6,11,13,15,26,27^. A family of recurrent ACE2-blocking hNAbs is composed of heavy chains (HCs) encoded by the *VH3-53* or *VH3-66* gene segment^3,12,13,16,17,27–29^, a majority of which are known or predicted^15,26,28,30,31^ to exhibit a common RBD binding mode resulting from the use of germline-encoded residues within the complementarity-determining regions 1 and 2 (CDRH1 and CDRH2) and a CDRH3 that is shorter than the average length (15 amino acids; IMGT^32^ CDR definition) in human antibodies^33^. Other SARS-CoV-2 RBD-binding antibodies are encoded by *VH3-30*^5^, which have also been isolated from SARS-CoV-infected donors^34^, and antibodies with a variety of the other VH gene segments^3,5,10,12–17^.

To classify commonalities and differences among RBD-binding hNAbs isolated from convalescent COVID-19 individuals^5^, we solved complexes of hNAbs with stabilized (2P and 6P versions)^35,36^ of soluble S trimer and used high-resolution details of the binding orientations of *VH1-2*, *VH1-46*, *VH3-30*, *VH3-53, VH4-34*, and *VH5-51* and hNAbs to elucidate rules for binding by four distinct anti-RBD antibody classes (Supplementary Table 2). The hNAbs chosen for structures are highly potent, achieving 90% neutralization in pseudotype virus assays at concentrations ranging from 22-140 ng/mL^5^, thus our structural analyses and classifications directly relate to understanding mechanisms of neutralization and potency differences between hNAbs.

## Class 1: *VH3-53*/short CDRH3 hNAbs that block ACE2 binding and bind “up” RBDs

We solved Fab and Fab-RBD crystal structures of C102 (Supplementary Table 1), which we compared to our previous cryo-EM structure of S trimer complexed with the related hNAb C105^26^ (Extended Data Fig. 1,2). Both C102 and C105 are *VH3-53* hNAbs with short (9 and 12 residues) CDRH3s (Extended Data Fig. 1g) that were isolated from the same donor^5^. They share structural similarities with each other and with other *VH3-53*/short CDRH3 hNAb structures solved as complexes with RBDs^12,30,37,38^ (Extended Data Fig. 2a). Importantly, the C102-RBD structure resembled the analogous portion of the C105-S structure^26^ (Extended Data Fig. 2a). These results establish that Fab-RBD structures can reproduce interactions with RBDs in the context of an S trimer; however, Fab-RBD structures do not reveal the state(s) of the antibody-bound RBD in the complex (“up” versus “down”) or the potential inter-protomer contacts by Fabs.

Since the C105 Fab bound either two or three “up” RBDs on S with no observed interactions with “down” RBDs or with adjacent RBDs^26^ (Extended Data Fig. 1f), we used the higher-resolution C102 Fab-RBD structure to deduce a more accurate epitope/paratope than possible using the C105-S cryo-EM structure with flexible “up” RBDs (Extended Data Fig. 1a-e). Buried surface area (BSA) calculations showed that the C102 CDRH3 played a relatively minor role in the paratope: of 1045 Å^2^ BSA on the antibody (786 Å^2^ on the HC; 259 Å^2^ on the light chain; LC), CDRH3 accounted for only 254 Å^2^ (Extended Data Fig. 2b). This contrasts with the majority of antibodies in which CDRH3 contributes equally or more to the interface with antigen than the sum of CDRH1 and CDRH2 contributions^39^. The epitopes on RBD for all available *VH3-53*/short CDRH3 hNAbs span the ACE2 binding site^15,26,28,30,31^ and show common RBD-binding interactions, represented by the C102 epitope (Extended Data Fig. 1b-e), which buried 1017 Å^2^ on RBD (Extended Data Fig. 2b). The ACE2-blocking epitope for these hNAbs is sterically occluded in the RBD “down” conformation (Fig. 1b; Extended Data Fig. 1f); therefore, class 1 hNAbs can only bind to “up” RBDs, as observed in the C105-S structure^26^, and as previously discussed, IgGs in this class could crosslink adjacent RBDs within a single trimer to achieve tighter binding through avidity effects^26^.

**Figure 1.**
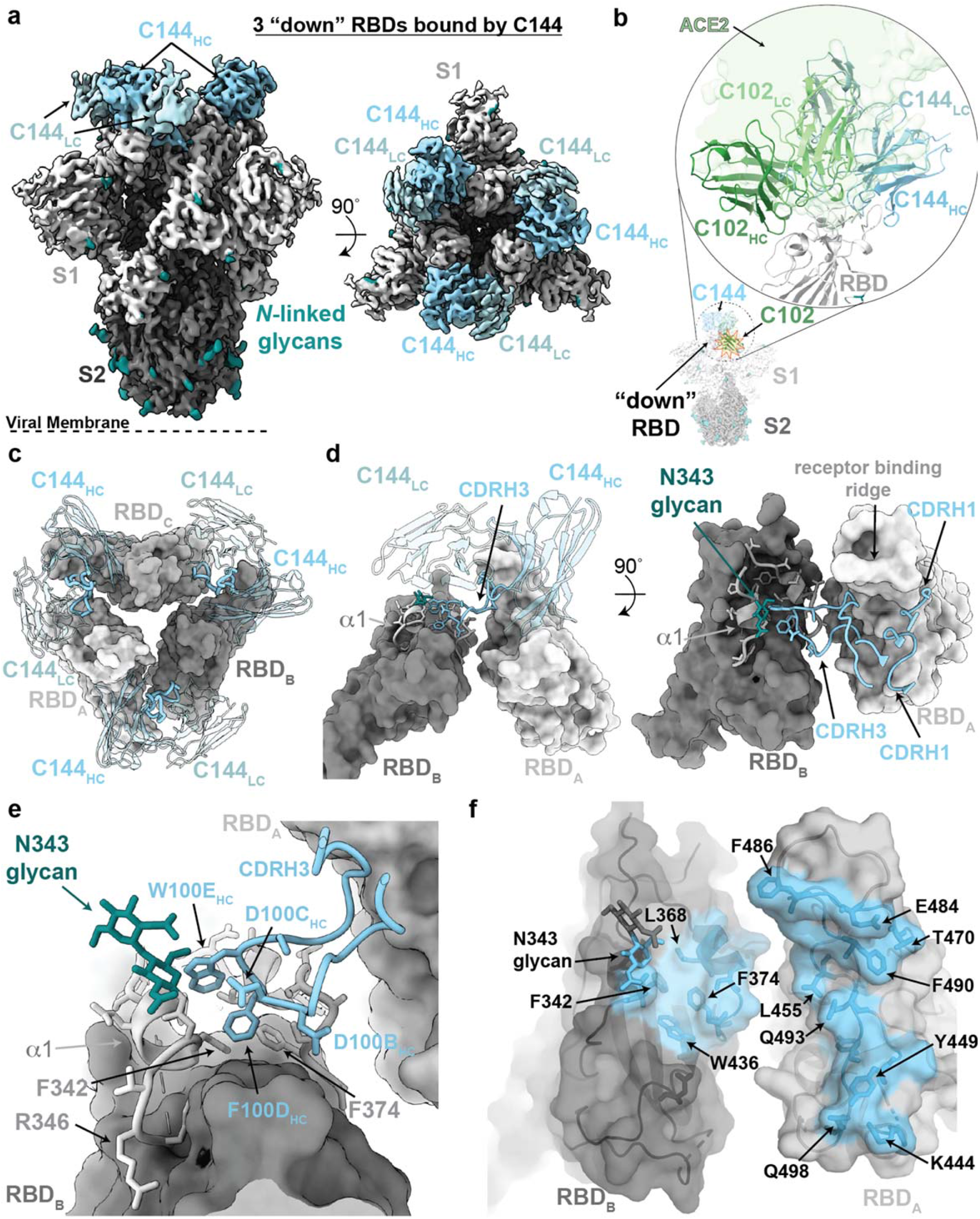
Cryo-EM structure of the C144-S complex illustrates a distinct *VH3-53* hNAb binding mode. **a,** 3.2 Å cryo-EM density for C144-S trimer complex revealing C144 binding to a closed (3 RBDs “down”) spike conformation. **b,** Overlay of C102 Fab (from C102-RBD crystal structure; Extended Data Fig. 1) and C144 Fab (from C144-S structure) aligned on a RBD monomer. ACE2 (PDB 6M0J; light green surface) is aligned on the same RBD for reference. C144 adopts a distinct conformation relative to the C102-like *VH3-53*/short CDRH3 NAb class, allowing binding to the “down” RBD conformation on trimeric spike, whereas C102-like NAbs can only bind “up” RBDs. **c,** Quaternary epitope of C144 involving bridging between adjacent RBDs via the CDRH3 loop. **d,e,** Close-up view of CDRH3-mediated contacts on adjacent protomer RBD (dark gray). C144 CDRH3 residues F100_D_ and W100_E_ are buried in a hydrophobic pocket comprising the RBD α1 helix, residue F374_RBD_ and the N343_RBD_-glycan. **f,** Surface representation of C144 epitope (light blue) across two adjacent RBDs. RBD epitope residues (defined as residues containing atom(s) within 4 Å of a Fab atom) are labeled in black.

## Class 2: hNAbs that overlap with the ACE2 binding site and recognize both “up” and “down” RBD conformations

In addition to the recurrent *VH3-53* hNAbs with short CDRH3s, a small subset of potently neutralizing *VH3-53* encoded antibodies utilize longer CDRH3s (>15 residues, IMGT definition^32^, Extended Data Fig. 1g)^5,12^. A recent structure of a RBD complexed with a *VH3-53*/long CDRH3 hNAb (COVA2-39) revealed a different RBD binding mode^38^, thus confirming predictions that binding with a C102-like interaction requires a short CDRH3^26,30^. To further elucidate molecular mechanisms for binding of *VH3-53*/long CDRH3 hNAbs, we solved a 3.2 Å cryo-EM structure of C144 (*VH3-53/VL2-14*; 25-residue CDRH3) bound to a S trimer^36^ (Extended Data Fig. 3). Despite the ability of ligand-free stabilized S trimers to adopt “up” RBD conformations^36^ and modeling suggesting the C144 binding site would be accessible on “up” RBDs (Fig. 1b), the C144-S structure revealed three C144 Fabs bound to a completely closed S with three “down” RBDs (Fig. 1a). The C144 binding mode differs from class 1 hNAbs, whose binding orientation is incompatible with “down” RBD conformations (Fig. 1b). In addition, the binding orientation observed for C144 differs from the binding described for COVA2-39, whose RBD epitope is predicted to be accessible only on “up” RBDs^38^ due to steric hinderances imposed on the LC by the N343_RBD_-associated glycan on the adjacent RBD (Extended Data Fig. 1h). Despite orientation differences, the RBD epitopes of C144, C102 and COVA2-39 overlap with the ACE2 binding site, suggesting a neutralization mechanism involving direct competition with ACE2 (Fig. 1b). ^40^

An interesting feature of C144 binding is that its long CDRH3 bridges between adjacent “down” RBDs to lock the spike glycoprotein into a closed, prefusion conformation, providing an additional neutralization mechanism in which S cannot open to engage ACE2 (Fig. 1c,d). The formation of C144’s quaternary epitope is driven by sandwiching CDRH3 residues F100_D_ and W100_E_ into a hydrophobic RBD cavity at the base of an *N*-linked glycan attached to N343_RBD_. The cavity comprises the RBD α1 helix (337-344), α2 helix (364-371), and hydrophobic residues (F374_RBD_ and W436_RBD_) at the edge of the RBD 5-stranded β-sheet (Fig. 1e,f). By contrast to CDRH3s of class 1 *VH3-53*/short CDRH3 hNAbs, C144’s CDRH3 contributed to a majority (∼60%) of the paratope and buried 330 Å^2^ surface area on the adjacent RBD (Extended Data Fig. 2b), likely explaining observed escape at residue L455_RBD_ (Fig. 1f) in C144 selection experiments^40^. Despite adjacent hydrophobic residues (F100_D_ and W100_E_) likely to be solvent-exposed before antigen binding, C144 IgG showed no evidence of non-specific binding in a polyreactivity assay (Extended Data Fig. 1i).

Given the unusual binding characteristics of C144, we investigated whether antibodies that showed similar S binding orientations in low-resolution negative-stain EM (nsEM) reconstructions^5^ utilize similar neutralization mechanisms. We characterized Fab-S cryo-EM structures (overall resolutions from 3.4-3.8 Å) of potent hNAbs (C002, C104, C119, and C121) predicted to compete with ACE2 binding^5^, which varied in their V gene segment usage and CDRH3 lengths (Fig. 2, Extended Data Figs. 3,4; Extended Data Table 1). Fab-S cryo-EM structures of these class 2 hNAbs showed bound RBDs in both “up” or “down” conformations, consistent with observations of similar hNAbs from nsEM^5,12^ and single-particle cryo-EM studies^10,34,41^. By contrast, the C144-S structure showed Fabs bound only to “down” RBDs (Fig. 1), suggesting that C144 binding requires recognition of the closed S trimer, or that C144 Fab(s) initially bound to “up” RBD(s) could trap the closed (3 RBDs “down”) S conformation through CDRH3-mediated interactions between adjacent RBDs.

**Figure 2.**
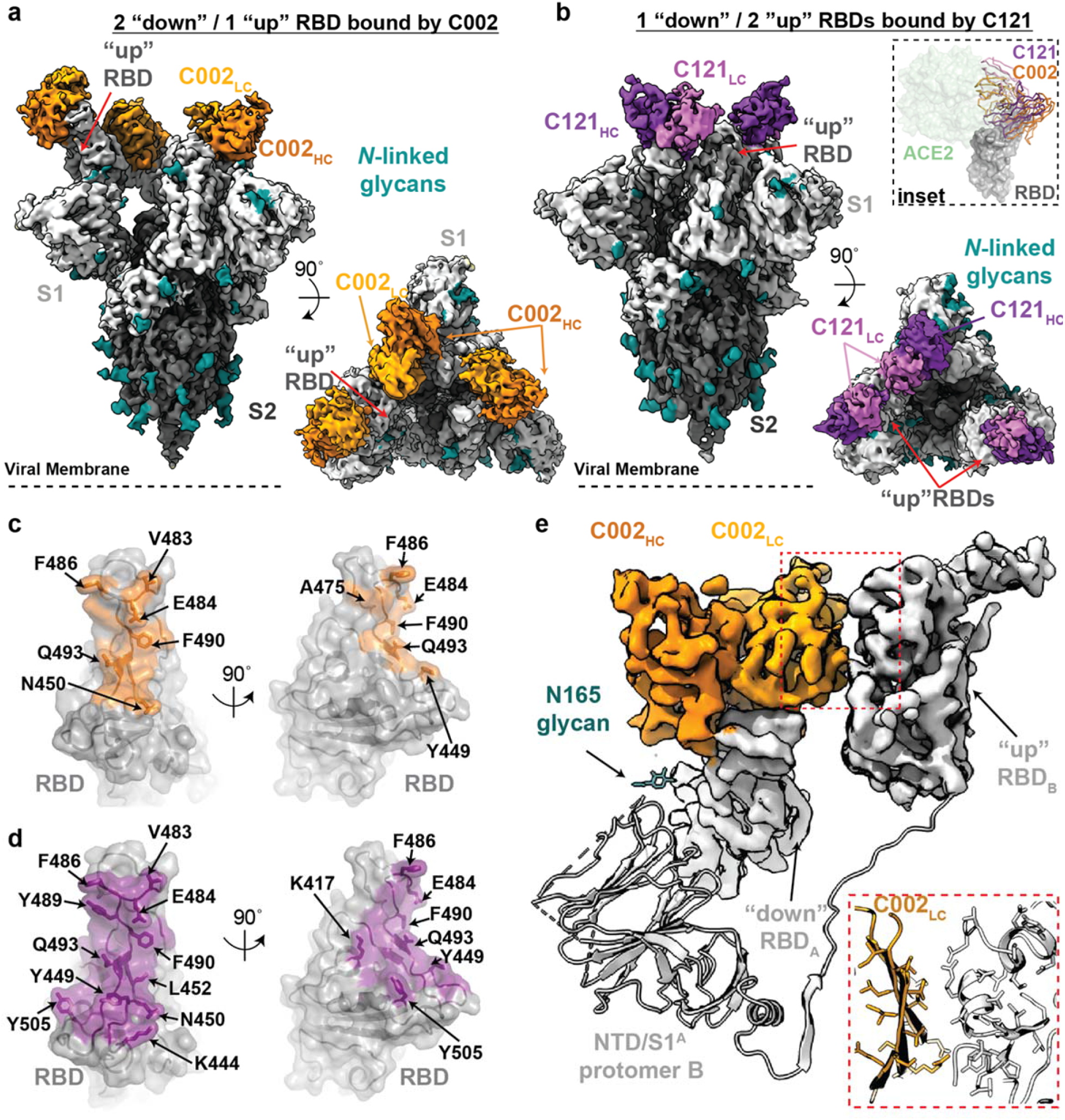
Cryo-EM structures of class 2 C002 and C121 hNAbs show binding to “up” and “down” RBDs. **a,b**, Cryo-EM densities for C002-S (panel a; 3.4 Å) and C121-S complexes (panel b; 3.7 Å) revealing binding of C002 or C121 to both “down” and “up” RBDs. Inset: Alignment of C002 and C121 Fabs on the same RBD. ACE2 is represented as a green surface for reference. **c,d**, Surface representations of C002 epitope (orange, panel c) and C121 epitope (purple, panel d) on the RBD surface (gray). RBD epitope residues (defined as residues containing atom(s) within 4 Å of a Fab atom) are labeled in black. **e,** C002 forms inter-protomer contacts via binding to an adjacent “up” RBD conformation on the surface of the trimer spike (also observed for class 2 C121-, C119-, and C104-S structures, see Extended Data Fig. 5). Red box: Close-up of adjacent “up” RBD and C002 LC interface.

To better understand commonalities of class 2 RBD epitopes, we further analyzed two additional potent hNAbs, C002 *(VH3-30*/*VK1-39*, 17-residue CDRH3, IC_50_=8.0 ng/mL^5^) and C121 (*VH1-2*/*VL2-23*, 23-residue CDRH3, IC_50_=6.7 ng/mL^5^), for which cryo-EM Fab-S structures were solved to 3.4 Å and 3.6 Å, respectively (Fig. 2a,b) using crystal structures of unbound C002 and C121 Fabs for fitting (Supplementary Table 1). The C002 and C121 RBD epitopes are focused on the receptor-binding ridge, overlapping with polar and hydrophobic residues along the flat face of the RBD responsible for ACE2 interactions (Fig. 2c-e). Similar to C144, hNAbs C002 and C121 buried most of the RBD epitope against HC CDR loops, with LC CDR loops engaging the receptor-binding ridge (Fig. 3). Interestingly, Fab-S structures of C002, C121, C119 and C104 revealed a quaternary epitope involving an adjacent RBD (Extended Data Figs. 3,4, 5a-c), albeit distinct from the quaternary binding of C144 (Fig. 1c-e). This C102/C121/C119/C104 type of secondary interaction was only observed when a Fab was bound to a “down” RBD and adjacent to an “up” RBD. The extent of the secondary interactions varied depending on the antibody pose (Extended Data Fig. 5a-c). Bridging interactions between adjacent “up” and “down” RBDs would not allow the two Fabs of a single IgG to bind simultaneously to an S trimer. However, this class of antibodies could support bivalent interactions between two adjacent “down” RBDs (Extended Data Fig. 5h, Extended Data Table 1).

**Figure 3.**
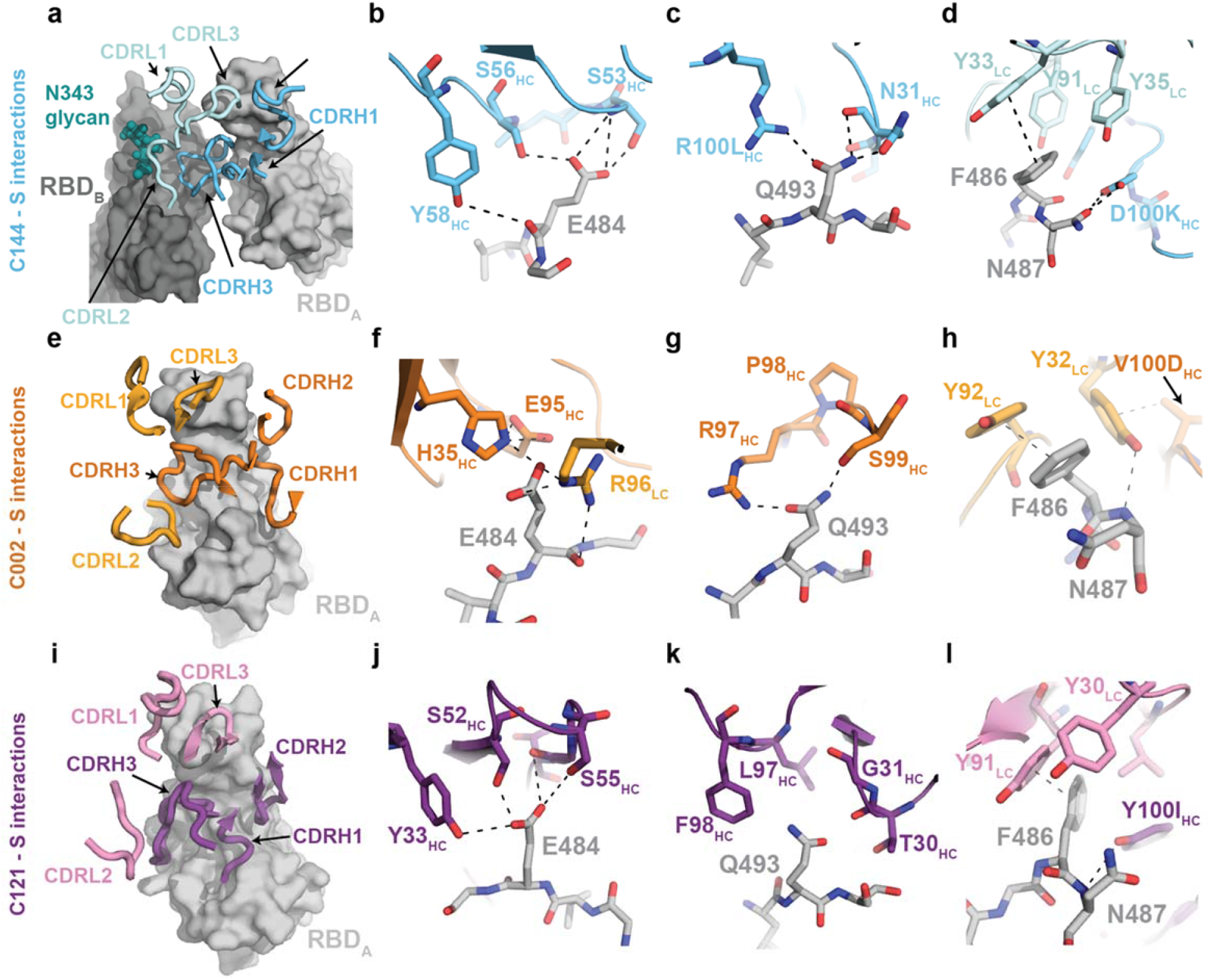
Details of common RBD interactions among class 2 hNAbs. Conserved interactions between the RBD and CDRs of class 2 NAbs as observed for **a-d,** C144 (HC: cyan, LC: sky blue), **e-h**, C002 (HC: dark orange, LC: light orange), and **i-l**, C121 (HC: purple, LC: pink). Primary and secondary epitopes on adjacent “down” RBDs are shown for C144. Secondary epitopes for C002 and C121, which require adjacent “up” RBDs, are shown in Extended Data Fig. 5. RBDs are gray; potential H-bonds and pi-pi stacking interactions (panel d, Y33_LC_ and F486_RBD_; panel h, Y92_LC_ and F486_RBD_; panel l, Y91_LC_ and F486_RBD_) are indicated by dashed lines.

Characterization of the highest resolution interface (C002-S structure) showed C002 LC framework regions (FWRs) 1 and 2 interfaced with the RBD residues comprising the 5-stranded β-sheet and α-helix that spans residues 440_RBD_–444_RBD_ (Fig. 2e), which is typically located near the three-fold axis of a closed S trimer. In addition to contacting neighboring RBDs, inter-protomer engagement with the N165_NTD_-associated glycan in the N-terminal domain (NTD) was observed for the class 2 hNAb BD23^13^. If fully processed, the N165_NTD_ glycan could adopt a conformation that would allow interactions with HC FWR3 and CDRH1 (Fig. 2e). However, in the structures reported here, we did not observe N165_RBD_ glycan density beyond the initial GlcNAc.

Given differences in class 2 hNAb V gene segments, CDRH3 lengths, and antibody poses, we investigated sequence features that drive conserved interactions. Sequence differences between SARS-CoV-2 and SARS-CoV RBD, including at positions 486_RBD_ and 493_RBD_ (F and Q, respectively, in SARS-CoV-2), in the ACE2 receptor-binding motif (RBM) allowed more favorable ACE2 binding to the SARS-CoV-2 RBD^42^. Analysis of interactions by C144, C002, and C121 revealed common interactions with these residues and also for E484_RBD_ by both antibody HC and LC residues (Fig. 3). In particular, class 2 hNAb interactions with F486_RBD_ mimicked ACE2 interactions, in that F486_RBD_ buries into a hydrophobic pocket typically involving CDRL1/CDRL3 tyrosine residues^43^ (Fig. 3d,h,l). Mimicking of the ACE2 F486_RBD_ binding pocket by SARS-CoV-2 hNAbs was observed across different LC V gene segments (Extended Data Table 1), suggesting that there is no restriction in LC V gene segment usage for class 2 hNAbs. Interestingly, a germline-encoded feature described for *VH3-53/*short CDRH3 class 1 hNAbs, the CDRH2 SxxS motif, is also found in other class 2 hNAbs (e.g., C121 and C119) despite different VH gene segment usage. Similar to *VH3-53* hNAbs C144 and COVA2-39, the C121 CDRH2 SxxS motif forms a potential hydrogen bond network with residue E484_RBD_ (Fig. 3b,j).

Overall, these results suggest a convergent mode of recognition by germline-encoded residues across diverse VH/VL gene segments for SARS-CoV-2, which may contribute to low levels of somatic hypermutation observed for these hNAbs (Extended Data Fig. 6, Extended Data Table 1).

## Class 3: hNAbs that bind outside the ACE2 binding site and recognize both “up” and “down” RBD conformations

C135 is a potent hNAb that showed binding properties distinct from class 1, class 2, and the cross-reactive SARS-CoV antibody CR3022^5^ (which we categorized as a class 4 antibody; Extended Data Table 1). To evaluate the mechanism of C135-mediated neutralization of SARS-CoV-2, we solved the cryo-EM structure of a C135-S complex to 3.5 Å (Fig. 4a, Extended Data Fig. 7), using an unbound C135 crystal structure for fitting (Supplementary Table 1). The structure revealed three C135 Fabs bound to an S trimer with 2 “down” and 1 “up” RBDs, although the C135-bound “up” RBD conformation was weakly resolved and therefore not modeled. C135 recognizes an glycopeptidic epitope similar to the cross-reactive SARS-CoV hNAb S309^34^, focusing on a region of the RBD near the N343_RBD_ glycan and non-overlapping with the ACE2 binding site (Fig. 4b, Extended Data Fig. 7c,d). Despite differences in binding orientations between C135 and S309, targeting of the RBD epitope was mainly V_H_-mediated (the BSA of RBD on the C135 HC represented ∼480Å^2^ of ∼700 Å^2^ total BSA) and included interactions with the core fucose moiety of the N343_RBD_ glycan. The smaller C135 footprint relative to S309 (∼700 Å^2^ versus ∼1150 Å^2^ BSA, respectively; Extended Data Fig. 7c,d) focused on interactions with RBD residues R346_RBD_ and N440_RBD_, which are engaged by residues from HC and LC CDR loops (Fig. 4c,d) and are not conserved between SARS-CoV-2 and SARS-CoV RBDs, rationalizing the lack of SARS-CoV cross-reactivity observed for C135^5^.

**Figure 4.**
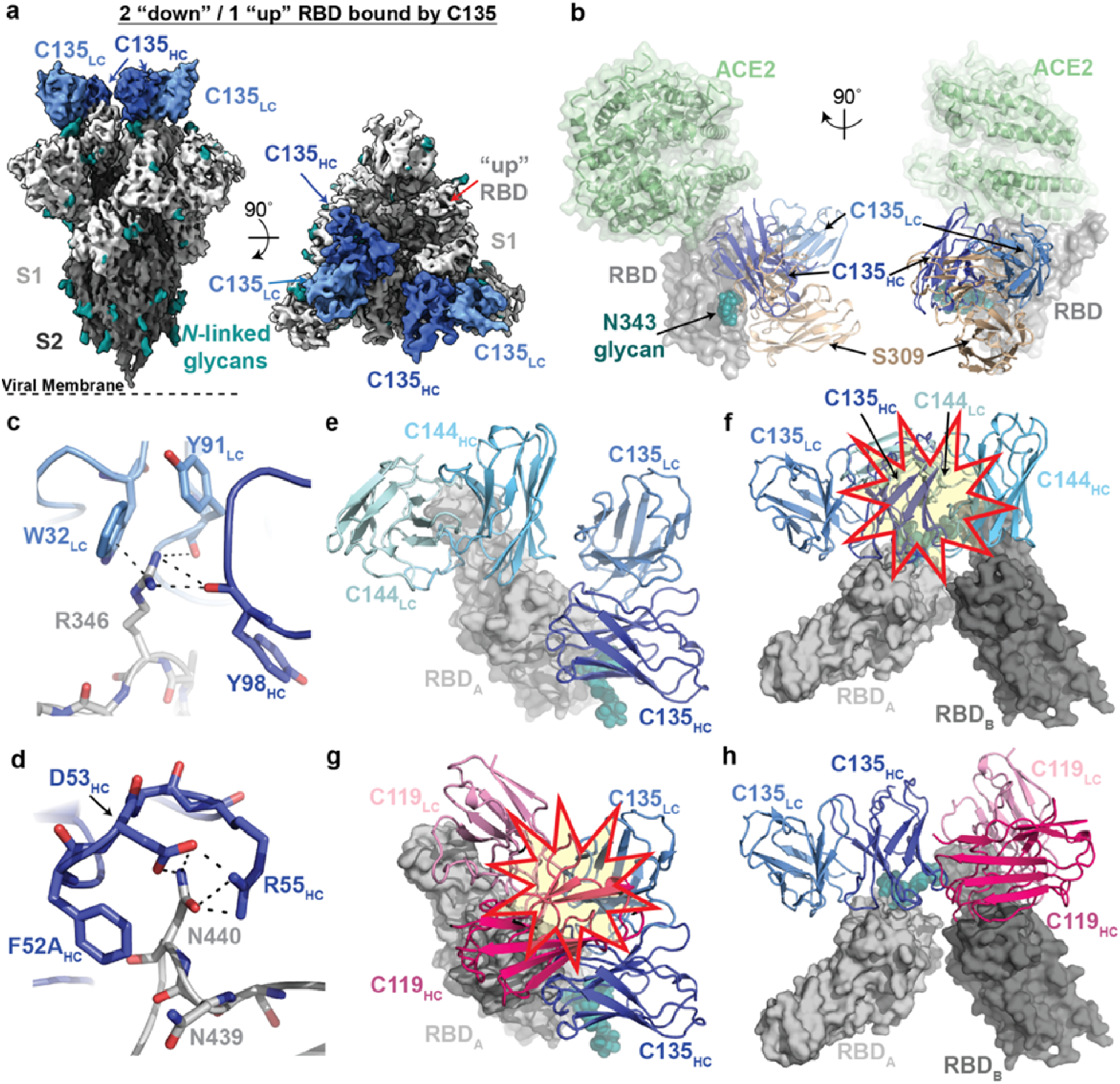
Cryo-EM structure of S complexed with the class 3 (non-ACE2 blocking) hNAb C135. **a**, 3.5 Å cryo-EM density of C135-S complex. **b,** Composite model of C135-RBD (blue and gray, respectively) overlaid with the SARS-CoV-2 NAb S309 (sand, PDB 6WPS) and soluble ACE2 (green, PDB 6M0J). The model was generated by aligning on 188 RBD Cα atoms. **c-d,** C135 CDRH (dark blue) and CDRL (light blue) interactions with residues R346_RBD_ (panel c) and N440_RBD_ (panel d). Potential pi-pi stacking interactions in c and H-bonds in c and d are illustrated by dashed black lines. **e-f,** Model of RBD interactions of NAbs C135 (class 3) and C144 (class 2) demonstrating that both Fabs can bind simultaneously to a single monomeric RBD (panel e), but would clash if bound to adjacent “down” RDBs on S trimer (panel f). Steric clashes indicated by a red and yellow star in f. **g-h,** Model of RBD interaction of NAbs C135 (class 3) and C119 (class 2) demonstrating that both Fabs cannot bind simultaneously to a single monomeric RBD (panel g), but do not clash if bound to adjacent “down” RDBs on S trimer (panel h). Steric clashes indicated by a red and yellow star in g.

The discovery of class 3 hNAbs such as C135 and S309 that were raised during SARS-CoV-2 or SARS-CoV natural infections, respectively, and bind outside of the ACE2 binding site, provides the potential for additive neutralization effects when combined with hNAbs that block ACE2, while also limiting viral escape^1,40^. A pair of antibodies in human clinical trials that includes REGN10987^8^, a hNAb that binds distal to the ACE2 binding site, prevented SARS-CoV-2 viral escape *in vitro*, but did not show synergistic neutralization^6^. Comparison of C135 and REGN10987 interactions with S showed similarities in epitopes (interactions focused on residues R346_RBD_ and N440_RBD_; Extended Fig. 7c,f). However, REGN10987 binding would sterically hinder ACE2 interactions, whereas C135 binding does not (Extended Data Fig. 7b, Fig. 4b). Interestingly, a structure of S complexed with C110 (*VH5-51*/*VK1-5*), isolated from the same donor as the C102 and C105 (class 1) and C119 and C121 (class 2) hNAbs^5^, showed a binding pose resembling REGN10987’s (Extended Data Fig. 7b,e-f). The C110 epitope showed similarities with both class 3 and class 2 hNAbs, binding distal to the ACE2 binding motif, but like REGN10987, could potentially sterically interfere with ACE2 (Extended Fig. 7). For each of these class 3 hNAbs, the Fab binding pose suggests that intra-protomer crosslinking by a single IgG is not possible (Extended Data Table 1).

Class 3 hNAbs add to the anti-SARS-CoV-2 antibody repertoire and could likely be effectively used in therapeutic combinations with class 1 or class 2 hNAbs. However, when using structures to predict whether hNAbs have overlapping epitopes, it is sometimes not sufficient to only examine Fab-RBD structures or even static images of S trimer because of the dynamic nature of the spike. Thus what might appear to be non-overlapping epitopes on an isolated RBD could overlap in some (Fig, 4e,f), but not all (Extended Data Fig. 8), scenarios on a spike trimer, complicating interpretation of competition experiments using monomeric RBDs and S trimers. The opposite can also be true; i.e., two Fabs that are predicted to be accommodated on a trimer could clash on an RBD monomer (Fig. 4g,h). Finally, adjacent monomers in different orientations could accommodate different antibodies that target overlapping sites (Extended Data Fig. 8).

## RBD substitutions affect hNAb binding to varying extents

VSV reporter viruses pseudotyped with SARS-CoV-2 S can escape by mutation from hNAbs C121, C135, or C144^40^, three of the antibodies used for the structural studies reported here. RBD mutations that were selected in response to antibody pressure correlated with the epitopes mapped from the structures of their Fabs complexed with S trimer (Fig. 1,2,4).

To further assess the effects of these and other RBD substitutions, we assayed hNAbs for which we obtained structural information (eight from this study; C105-S complex from ref.^26^) for binding to mutated RBD proteins. The RBD mutants included two that induced escape from the class 3 hNAb C135 (R346S and N440K)^40^ (Fig. 4c,d), one found in circulating isolates^44^ that conferred partial resistance to C135 (N439K)^40^ (Fig. 4d), a circulating variant (A475V) that conferred resistance to class 1 and 2 *VH3-53* hNAbs^44^, two that induced escape from C121 or C144 (E484K and Q493R)^40^ (Fig. 3), and a circulating variant that conferred partial resistance to C121 (V483A)^40^. Kinetic and equilibrium constants for the original and mutant RBDs were derived from surface plasmon resonance (SPR) binding assays in which RBDs were injected over immobilized IgGs (Extended Data Fig. 9). Loss of binding affinity was consistent with RBD mutations that conferred escape, with hNAbs within each class being similarly affected by the same point mutations, which was not seen when comparing effects of point mutations between hNAb classes. This suggests that antibody pressure that leads to escape from one hNAb class would be unlikely to affect a different class. These results suggest a therapeutic strategy involving hNAbs of different classes for monoclonal NAb treatment of SARS-CoV-2–infected individuals.

## Conclusions

The Fab-S structures reported here represent a comprehensive structural, biophysical, and bioinformatics analysis of SARS-CoV-2 NAbs (Extended Data Fig. 10), providing critical information for interpreting correlates of protection for clinical use. The structures reveal a wealth of unexpected interactions of hNAbs with the spike trimer, including five antibodies that reach between adjacent RBDs on the protomers of a single spike trimer. A dramatic example of bridging between spike protomers involved a hNAb, C144, that uses a long CDRH3 with a hydrophobic tip to reach across to an adjacent RBD, resulting in all three RBDs on spike trimer being locked into a closed conformation. This example, and the four other hNAbs that contact adjacent RBDs, demonstrates that crystal structures of Fab-monomeric RBD complexes, while informative for defining a primary epitope on one RBD, do not reveal how antibodies actually recognize the flexible “up”/”down” RBD conformations on the spike trimer that are targeted for neutralization on the virus. Indeed, our cryo-EM structures of Fab-spike trimer complexes showed many possible combinations of recognized RBDs: three “up,” two “up” and one “down,” one “up” and two “down,” and three “down,” with some structures showing three Fabs bound per trimer and others showing two Fabs bound per trimer. By analyzing the approach angles of antibodies bound to RBDs on spike trimers, we can predict whether a particular IgG can bind to a single spike trimer to gain potency through avidity effects, which would also render the antibody more resistant to spike mutations. In addition, structural information allowed us to assess RBD mutants that arose in circulating viral isolates and/or were obtained by *in vitro* selection. Taken together, this comprehensive study provides a blueprint for designing antibody cocktails for therapeutics and potential spike-based immunogens for vaccines.

## Methods

### Protein Expression

Expression and purification of SARS-CoV-2 ectodomains were conducted as previously described^26^. Briefly, constructs encoded the SARS-CoV-2 S ectodomain (residues 16-1206 of the early SARS-CoV-2 GenBank MN985325.1 sequence isolate with 2P^35^ or 6P^36^ stabilizing mutations, a mutated furin cleavage site between S1 and S2, a C-terminal TEV site, foldon trimerization motif, octa-His tag, and AviTag) were used to express soluble SARS-CoV-2 S ectodomains. Constructs encoding the SARS-CoV-2 RBD from GenBank MN985325.1 (residues 331-524 with C-terminal octa-His tag and AviTag) and mutant RBDs were made as described^26^, SARS-CoV-2 2P S, 6P S, and RBD proteins were purified from the supernatants of transiently-transfected Expi293F cells (Gibco) by nickel affinity and size-exclusion chromatography^26^. Peak fractions were identified by SDS-PAGE, and fractions corresponding to S trimers or monomeric RBDs were pooled and stored at 4°C. Fabs and IgGs were expressed, purified, and stored as described^45,46^.

### X-ray crystallography

Crystallization trials were carried out at room temperature using the sitting drop vapor diffusion method by mixing equal volumes of a Fab or Fab-RBD complex and reservoir using a TTP LabTech Mosquito robot and commercially-available screens (Hampton Research). Crystals were obtained in 0.2 M ammonium sulfate, 20% w/v PEG 3350 (C102 Fab), 0.2 M sodium citrate tribasic, 20% w/v PEG 3350 (C102-RBD), 0.2 M lithium sulfate monohydrate, 20% w/v PEG 3350 (C002 Fab), 0.04 M potassium phosphate, 16% w/v PEG 8000, 20% v/v glycerol (C135 Fab), 0.2 M ammonium citrate pH 5.1, 20% PEG 3350 (C121 Fab), or 0.2 M sodium tartrate dibasic dihydrate pH 7.3, 20 % w/v PEG 3350 (C110 Fab). A C135 Fab crystal was directly looped and cryopreserved in liquid nitrogen. Other crystals were quickly cryoprotected in a mixture of well solution with 20% glycerol and then cryopreserved in liquid nitrogen.

X-ray diffraction data were collected for Fabs and the Fab-RBD complex at the Stanford Synchrotron Radiation Lightsource (SSRL) beamline 12-1 on a Pilatus 6M pixel detector (Dectris) at a wavelength of 1.0 Å. Data from single crystals of C121 Fab and C110 Fab were indexed and integrated in XDS^47^ and merged using AIMLESS in *CCP4*^48^ (Supplementary Table 1). Data from single crystals of C102 Fab, C135 Fab, and C002 fab were indexed and integrated using XDS^47^ and merged in Phenix^49^. Diffraction data for C002 Fab were anisotropically truncated and scaled using the UCLA Anisotropy Server^50^ prior to merging. Data from a single crystal of C102 Fab-RBD complex were indexed and integrated using XIA2^51^ implementing DIALS^52,53^ and merged using AIMLESS in *CCP4*^48^. For C110 Fab and C121 Fabs, structures were determined by molecular replacement in PHASER^54^ using the coordinates for B38 (PDB 7BZ5) or an inferred germline form of the HIV-1 NAb IOMA^55^ inferred germline (unpublished), respectively, after removing CDR loops as a search model. For C002 Fab, C102 Fab, C102 Fab-RBD, and C135 Fab, structures were determined by molecular replacement in PHASER^54^ using B38 Fab coordinates (PDB 7BZ5) after trimming HC and LC variable domains using Sculptor^56^ (and for the C102 Fab-RBD data, also RBD coordinates from PDB 7BZ5) as search models. Coordinates were refined using Phenix^49^ and cycles of manual building in Coot^57^ (Supplementary Table 1).

### Cryo-EM Sample Preparation

Purified Fabs were mixed with SARS-CoV-2 S 2P trimer^35^ or SARS-CoV-2 S 6P trimer^36^ (1.1:1 molar ratio Fab per protomer) to a final Fab-S complex concentration of 2-3 mg/mL and incubated on ice for 30 minutes. Immediately before deposition of 3 μL of complex onto a 300 mesh, 1.2/1.3 AuUltraFoil grid (Electron Microscopy Sciences) that had been freshly glow-discharged for 1 min at 20 mA using a PELCO easiGLOW (Ted Pella), a 0.5% w/v octyl-maltoside, fluorinated solution (Anatrace) was added to each sample to a final concentration of 0.02%. Samples were vitrified in 100% liquid ethane using a Mark IV Vitrobot (Thermo Fisher) after blotting at 22°C and 100% humidity for 3 s with Whatman No. 1 filter paper.

### Cryo-EM Data Collection and Processing

Single-particle cryo-EM data were collected on a Titan Krios transmission electron microscope (Thermo Fisher) operating at 300 kV for all Fab-S complexes except for C144-S, which was collected on a Talos Arctica (Thermo Fisher) operating at 200 kV. Movies were collected using SerialEM automated data collection software^58^ with beam-image shift over a 3 by 3 pattern of 1.2 µm holes with 1 exposure per hole. Movies were recorded in super-resolution mode on a K3 camera (Gatan) for the C144-S dataset on the Arctica (0.435 Å/pixel) or on a K3 behind BioQuantum energy filter (Gatan) with a 20 eV slit on the Krios (0.418 Å/pixel) for all other datasets. Data collections parameters are summarized in Supplementary Table 2. In general, the data processing workflow described below was performed for all data sets in cryoSPARC v2.15^59^.

Cryo-EM movies were patch motion corrected for beam-induced motion including dose weighting within cryoSPARC^59^ after binning super-resolution movies. The non-dose-weighted images were used to estimate CTF parameters using CTFFIND4^60^ or with cryoSPARC implementation of the Patch CTF job, and micrographs with power spectra that showed poor CTF fits or signs of crystalline ice were discarded. A subset of images were randomly selected and used for reference-free particle picking using Blob picker in cryoSPARC^59^. Particles were subjected to 2D classification and the best class averages that represented different views were used to generate 3 *ab initio* models. The particles from the best classes were used in another 2D classification job, and the best set of unique views was utilized as templates for particle picking on the full set of images. Initial particle stacks were extracted, down-sampled x2, and used in heterogeneous refinement against the 3 *ab initio* volumes generated with the smaller dataset (*ab initio* volumes used were interpreted as a Fab-S complex, free Fab or dissociated S protomers, and junk/noise class). Particles assigned to the Fab-S volume were further cleaned via iterative rounds of 2D classification to select class averages that displayed unique views and secondary structural elements. Resulting particle stacks were homogenously refined before being split into 9 individual exposure groups based upon collection holes. Per particle CTF and aberration corrections were performed and the resulting particles further 3D refined. Additional processing details are summarized in Supplementary Table 2.

Given the known heterogeneity of spike trimers^20,21^, homogenously refined particles were used for 3D classification in cryoSPARC^59^ (*ab initio* job: k=4 classes, class similarity=0.3). This typically resulted in one or two majority Fab-S complexes, with the other minority populated classes representing junk or unbound S trimer. Particles from the good class(es) were further subjected to 3D classification (*ab initio* job: k=4, class similarity=0.7) to attempt to separate various Fab-S complex states. If multiple states were identified (as observed for C002-S and C121-S complexes), particles were heterogeneously refined, followed by re-extraction without binning (0.836Å/pixel) before homogeneous refinement of individual states. For all other datasets, the majority of particles represented one state that was homogenously refined after re-extraction without binning.

Particle stacks for individual states were non-uniform refined with C1 symmetry and a dynamic mask. To improve resolution at the Fab-RBD interfaces, volumes were segmented in Chimera^61^ and the regions corresponding to the NTD_S1_/RBD_S1_ domains and Fab V_H_-V_L_ domains were extracted and used to generate a soft mask (5-pixel extension, 10-pixel soft cosine edge). Local refinements with the mask resulted in modest improvements of the Fab-RBD interface, which allowed for fitting and refinement of this region. The particles were then subjected to CTF refinement and aberration correction, followed by a focused, non-uniform refinement with polished particles imposing C1 symmetry (except for the C144-S complex where C3 symmetry was utilized). Final overall resolutions were according to the gold-standard FSC^62^. Details of overall resolution and locally-refined resolutions according to the gold-standard FSC^62^ can be found in Supplementary Table 2.

### Cryo-EM Structure Modeling and Refinement

Coordinates for initial complexes were generated by docking individual chains from reference structures into cryo-EM density using UCSF Chimera^63^. The following coordinates were used: SARS-CoV-2 S trimers: PDBs 6VYB and 6XKL, “up” RBD conformations: PDB 7BZ5, unbound C102, C002, C110, C135 Fab structures (this study) (Supplementary Table 1). Initial models were then refined into cryo-EM maps using one round of rigid body refinement followed by real space refinement. Sequence-updated models were built manually in Coot^57^ and then refined using iterative rounds of refinement in Coot^57^ and Phenix^49^. Glycans were modeled at potential *N*-linked glycosylation sites (PNGSs) in Coot^57^ using ‘blurred’ maps processed with a variety of B-factors^64^. Validation of model coordinates was performed using MolProbity^65^ (Supplementary Table 2).

### Structural Analyses

CDR lengths were calculated based on IMGT definitions^32^. Structure figures were made with PyMOL (Version 1.8.2.1 Schrodinger, LLC) or UCSF ChimeraX^61^. Local resolution maps were calculated using cryoSPARC v 2.15^59^. Buried surface areas were calculated using PDBePISA^66^ and a 1.4 Å probe. Potential hydrogen bonds were assigned as interactions that were <4.0Å and with A-D-H angle >90°. Potential van der Waals interactions between atoms were assigned as interactions that were <4.0Å. Hydrogen bond and van der Waals interaction assignments are tentative due to resolution limitations. RMSD calculations following pairwise Cα alignments were done in PyMOL without rejecting outliers. Criteria for epitope assignments are described in figure legends.

To evaluate whether intra-spike crosslinking by an IgG binding to a single spike trimer was possible (Extended Data Table 1), we first measured the Cα distance between a pair of residues near the C-termini of adjacent Fab C_H_1 domains (residue 222_HC_ on each Fab) (Extended Data Fig. 5h). We compared this distance to the analogous distances in crystal structures of intact IgGs (42 Å, PDB 1HZH; 48 Å, PDB 1IGY; 52 Å, PDB 1IGT). To account for potential influences of crystal packing in these measurements, as well as flexibility in the V_H_-V_L_/C_H_1-C_L_ elbow bend angle and uncertainties in C_H_1-C_L_ domain placement in Fab-S cryo-EM structures, we set a cut-off of ≤65 Å for this measured distance as possibly allowing for a single IgG to include both Fabs. Entries in the “Potential IgG intra-spike binding” column in Extended Data Table 1 are marked “No” if all of the adjacent Fabs in cryo-EM classes of that structure are separated by >65 Å for this measured distance. Entries in the “Potential IgG intra-spike binding” column in Extended Data Table 1 are marked as “Yes” if at least one pair of the adjacent Fabs in cryo-EM classes of that structure are separated by ≤65 Å for this measured distance.

### Surface plasmon resonance (SPR) binding experiments

SPR experiments were performed using a Biacore T200 instrument (GE Healthcare). IgGs were immobilized on a CM5 chip by primary amine chemistry (Biacore manual) to a final response level of ∼3000 resonance units (RUs). Concentration series of the original SARS-Cov-2 RBD and RBD mutants (six 4-fold dilutions starting from a top concentration of 1000 nM) were injected at a flow rate of at a flow rate of 30 μL/min over immobilized IgGs for a contact time of 60 sec, followed by a injection of 0.01 M HEPES pH 7.4, 0.15 M NaCl, 3 mM EDTA, 0.005% v/v surfactant P20 buffer for a dissociation time of 300 sec. Binding reactions were allowed to reach equilibrium, and *K*_D_s were calculated from the ratio of association and dissociation rates (*K*_D_ = *k*_d_/*k*_a_) derived from a 1:1 binding model (C002, C102, C105, C110, and C119 (except for C119-E484K), C121, C135, and C144), or from a two-state binding model (*K*_D_ = *k*_d_1/*k*_a_1 × *k*_d_2/[k_d_2+*k*_a_2]) (C104, C119-E484K). Kinetic constants were calculated using Biacore T200 Evaluation Software v3.2 using a global fit to all curves in each data set. Flow cells were regenerated with 10 mM glycine pH 2.0 at a flow rate of 90 μL/min.

### Polyreactivity assays

IgGs were evaluated for off-target interactions by measuring binding to baculovirus extracts containing non-specific proteins and lipids as described^60^. The assays were automated on a Tecan Evo2 liquid handling robot fitted with a Tecan Infinite M1000 plate reader capable of reading luminescence. Maxisorb 384-well plates (Nunc) were adsorbed overnight with a 1% preparation of recombinant baculovirus particles generated in Sf9 insect cells^67^. The adsorbed plate was blocked with 0.5% BSA in PBS, then incubated with 20 µL of a 1.0 µg/mL solution of IgG in PBS for 3 hours. Polyreactivity was quantified by detecting bound IgG using an HRP-conjugated anti-human IgG secondary antibody (Genscript) and SuperSignal ELISA Femto Maxiumum Sensitivity Substrate (Thermo Scientific). Relative Light Units (RLU) were measured at 475 nm in the integrated plate reader. Engineered human anti-HIV-1 IgGs previously demonstrated to exhibit high levels of polyreactivity (NIH45-46^G54W^ and 45-46m2)^61,62^ were used as positive controls. NIH45-46, which exhibited intermediate polyreactivity^63^, was also evaluated for comparisons. Negative control IgGs with low polyreactivity included the human HIV-1 antibodies N6^64^ and 3BNC117^63^ and bovine serum albumin (BSA). RLU values are presented as the mean and standard deviation of triplicate measurements in Extended Data Fig. 1i.

### Reporting Summary

Further information on research design is available in the Nature Research Reporting Summary linked to this paper.

## Data availability

The cryo-EM maps and atomic models will be deposited at the EMDB and the PDB. Crystal structure data will be deposited in the PDB. Described materials will be available upon request, in some cases after completion of a materials transfer agreement.

## Acknowledgements

We thank Dr. Jost Vielmetter, Pauline Hoffman, and the Protein Expression Center in the Beckman Institute at Caltech for expression assistance, Drs. Jost Vielmetter and Jennifer Keeffe for setting up automated polyreactivity assays, Dr. Jennifer Keeffe for construct design, and Nicholas Koranda for help with cloning and protein purification. Electron microscopy was performed in the Caltech Beckman Institute Resource Center for Transmission Electron Microscopy with assistance from Dr. Songye Chen. We thank the Gordon and Betty Moore and Beckman Foundations for gifts to Caltech to support the Molecular Observatory (Dr. Jens Kaiser, director), and Drs. Silvia Russi, Aina Cohen, and Clyde Smith and the beamline staff at SSRL for data collection assistance. Use of the Stanford Synchrotron Radiation Lightsource, SLAC National Accelerator Laboratory, is supported by the U.S. Department of Energy, Office of Science, Office of Basic Energy Sciences under Contract No. DE-AC02-c76SF00515. The SSRL Structural Molecular Biology Program is supported by the DOE Office of Biological and Environmental Research, and by the National Institutes of Health, National Institute of General Medical Sciences (P41GM103393). The contents of this publication are solely the responsibility of the authors and do not necessarily represent the official views of NIGMS or NIH. This work was supported by NIH grant P01-AI138938-S1 (P.J.B. and M.C.N.), the Caltech Merkin Institute for Translational Research (P.J.B.), NIH grant P50 8 P50 AI150464-13 (P.J.B.), and a George Mason University Fast Grant (P.J.B.). C.O.B was supported by the Hanna Gray Fellowship Program from the Howard Hughes Medical Institute and the Postdoctoral Enrichment Program from the Burroughs Wellcome Fund. M.C.N. is a Howard Hughes Medical Institute Investigator.

## Author contributions

C.O.B., M.C.N., A.P.W., and P.J.B. conceived the study and analyzed data; D.F.R. and M.C.N. provided monoclonal antibody sequences and plasmids derived from COVID-19 convalescent donors. C.O.B. and K.H.T. performed protein purifications and C.O.B. assembled complexes for cryo-EM and X-ray crystallography studies. C.O.B. performed cryo-EM and interpreted structures with assistance from M.A.E., K.A.D, S.R.E., A.G.M., and N.G.S. C.A.J. and C.O.B. performed and analyzed crystallographic structures, with refinement assistance from M.A.E and K.M.D. Y.E.L. performed polyreactivity assays. H.B.G. performed and analyzed SPR experiments. A.P.W. analyzed antibody sequences. C.O.B., M.C.N., A.P.W., and P.J.B. wrote the paper with contributions from other authors.

---

**Extended Data Figure 1:**
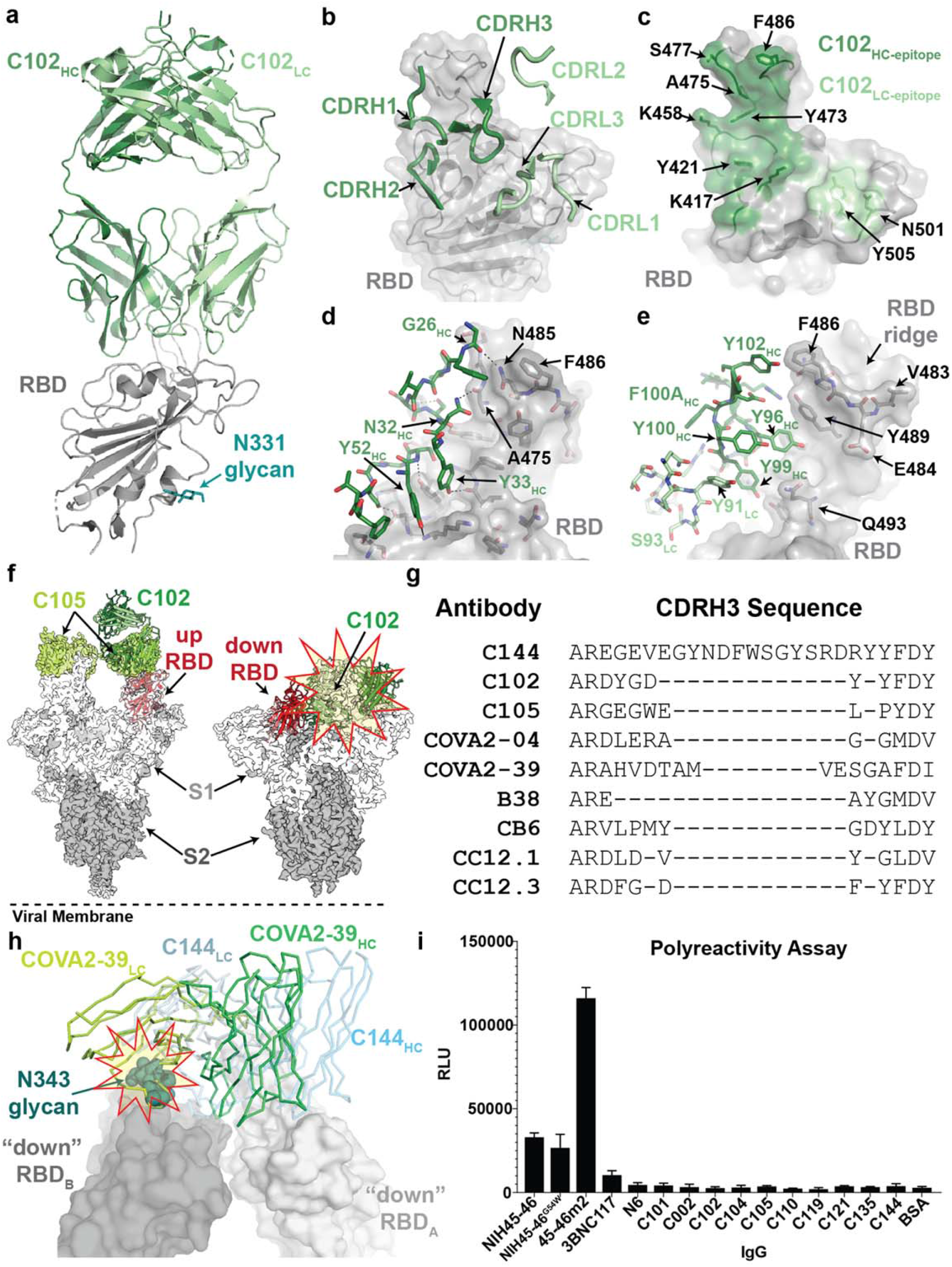
X-ray structure and epitope mapping of *VH3-53* hNAb C102. **a,** X-ray structure of C102 Fab – RBD_331-518_ complex. **b,** C102 CDR loops mapped on the RBD surface. **b,** Surface representation of C102 epitope colored by C102 HC (dark green) and LC (light green) interactions. **c,** CDRH1, CDRH2 and **d,** CDRH3 interactions with RBD residues. Potential H-bond contacts are illustrated as dashed lines. **f**, Left: Overlay of C102-RBD crystal structure (cartoon) with C105-S trimer cryoEM density (PDB 6XCM, EMD-22127) illustrating conserved binding to RBD epitope in an “up” conformation. Right: The C102 epitope is sterically occluded when aligned to a “down” RBD conformation (red and yellow star). SARS-CoV-2 S domains are dark gray (S2 domain) and light gray (S1 domain); the C105 Fab is yellow-green. **g,** Alignment of selected CDRH3 sequences for *VH3-53/VH3-66* SARS-CoV-2 neutralizing antibodies (IMGT definition^15^). **h**, Overlay of hNAb COVA2-39 Fab^4^ (lime green and lemon, from COVA2-39-RBD structure, PDB 7JMP) and C144 Fab (blue, from C144-S structure) aligned on a RBD_A_ of C144 epitope. COVA2-39 adopts a distinct conformation relative to the C102-like *VH3-53*/short CDRH3 NAb class and to C144, recognizing its RBD epitope only in an “up” RBD conformations due to steric clashes (red and yellow star) with the N343_RBD_-associated glycan on the adjacent RBD. **i,** Polyreactivity assay. IgGs were evaluated for binding to baculovirus extracts to assess non-specific binding. Polyreactive positive control IgGs were NIH45-46, NIH45-46^G54W^, and 45-46m2. Negative controls were bovine serum albuminn (BSA) and IgGs N6 and 3BNC117. Relative Light Unit (RLU) values are presented as the mean and standard deviation of triplicate measurements.

**Extended Data Figure 2.**
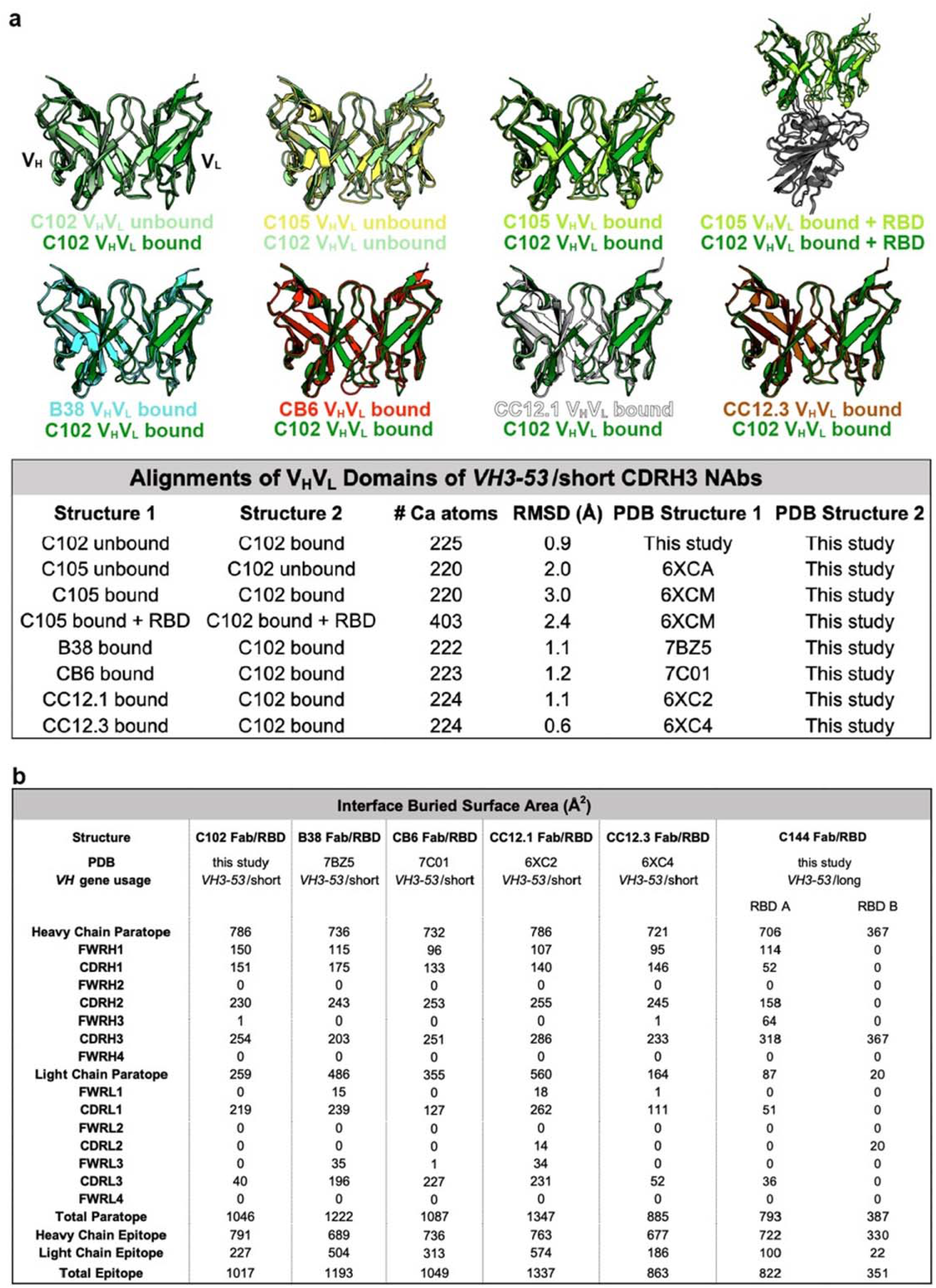
Overview of *VH3-53*/*VH3-66* hNAb structures. **a,** Superimposition of V_H_ and V_L_ domains of C102 with other *VH3-53*/*VH3-66* NAbs (top) and RMSD calculations (bottom). **b**, BSA comparisons for the indicated Fab/RBD structures. BSAs were calculated using PDBePISA^16^ and a 1.4 Å probe.

**Extended Data Figure 3.**
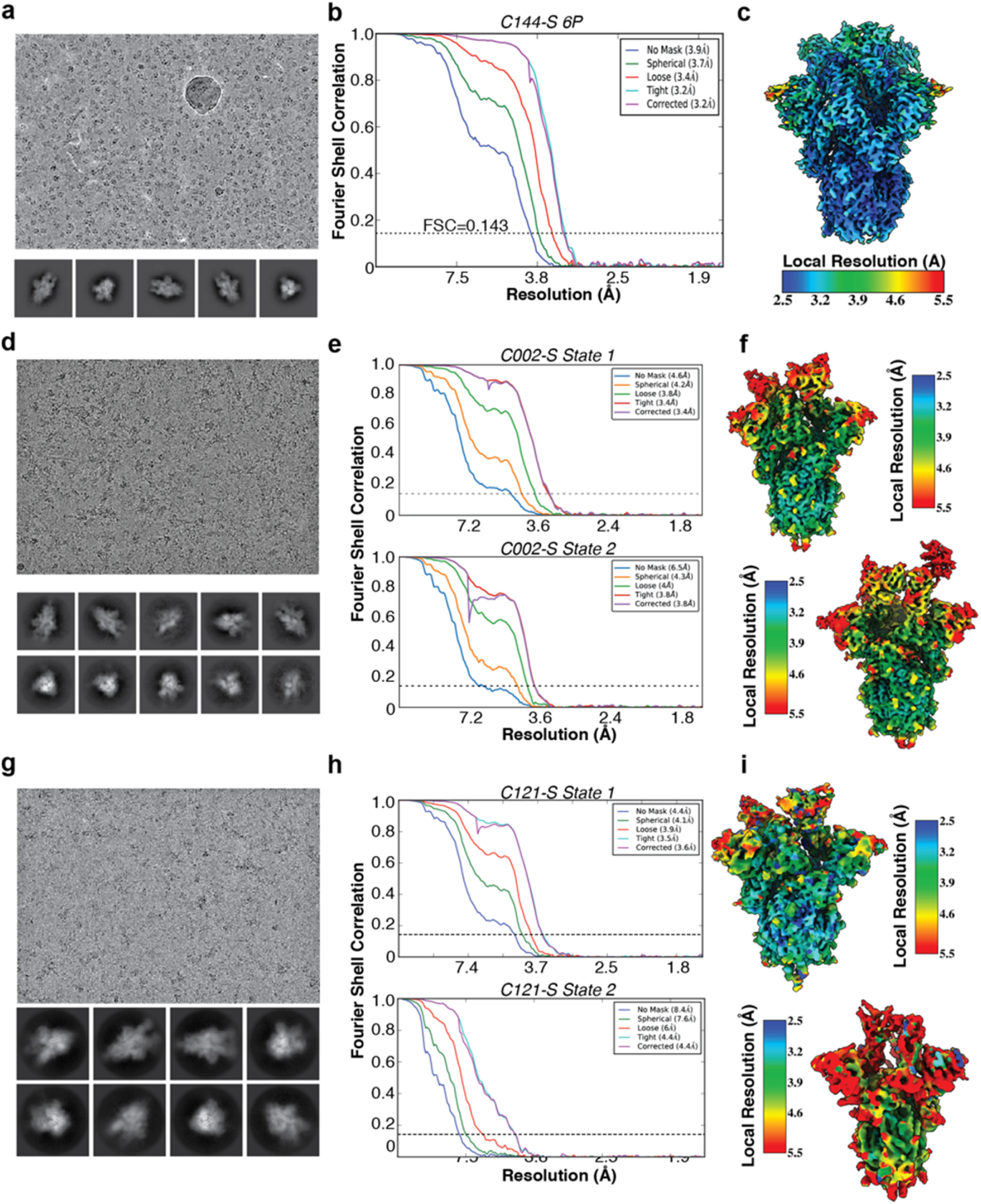
Cryo-EM data processing and validation for C144-S, C002-S, and C121-S complexes. Representative micrograph, 2D class averages, gold-standard FSC plots, and local resolution estimations for **a-c,** C144-S 6P, **d-f**, C002-S 2P, and **g-I**, C121-S 2P. For the C002-S dataset, two classes were resolved: State 1, C002 Fabs bound to 3 “down” RBDs, and State 2, C002 Fabs bound to 2 “down”/1 “up” RBD. For the C121-S 2P dataset, two classes were resolved: State 1, C121 Fabs bound to 2 “down”/1 “up” RBD and State 2, C121 Fabs bound to 1 “down”/2 “up” RBDs.

**Extended Data Figure 4.**
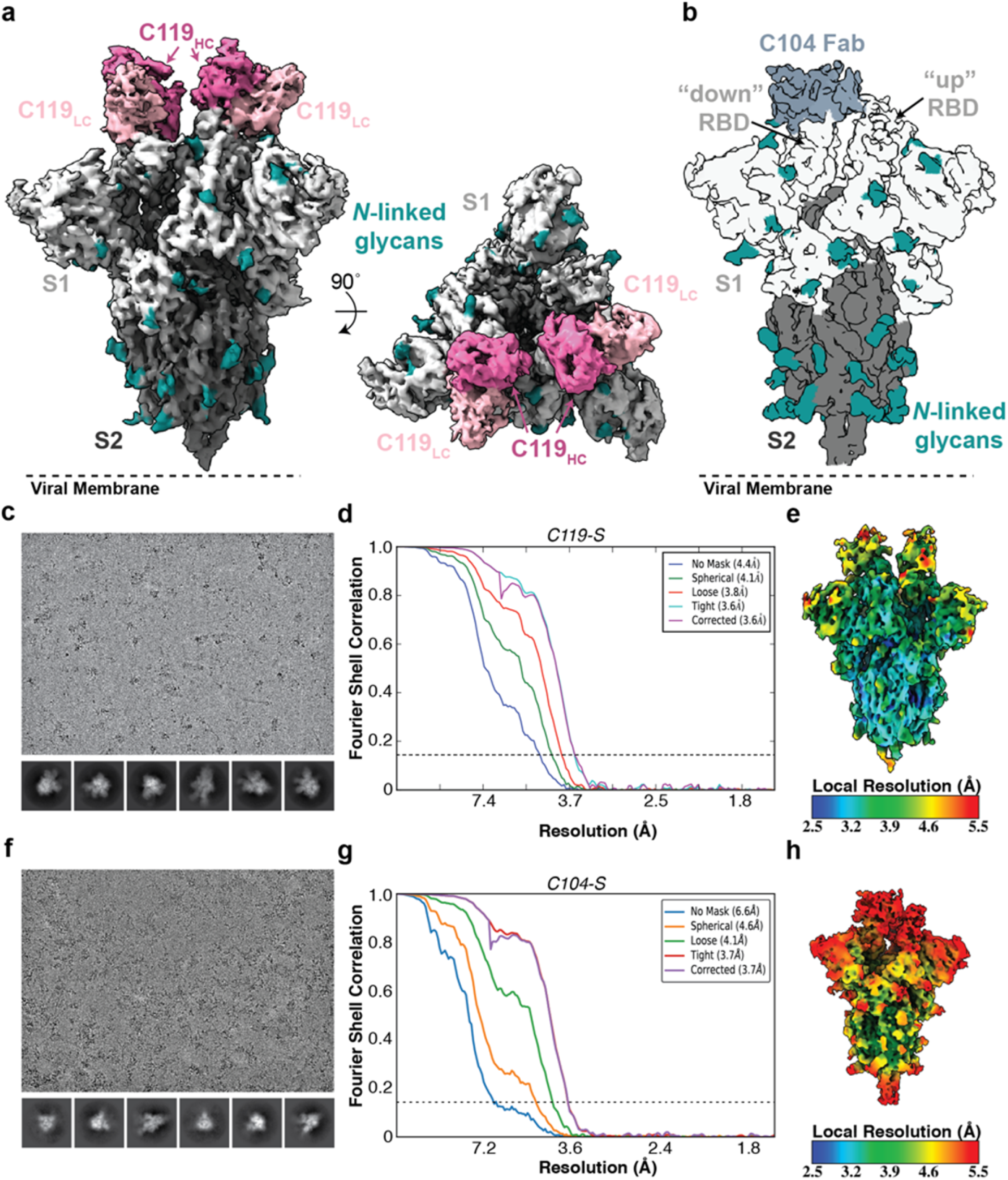
Cryo-EM processing, validation, and reconstruction for C119-S and C104-S complexes. **a,** 3.6 Å cryo-EM reconstruction for a C119-S trimer complex. **b,** 3.7 Å cryo-EM reconstruction for a C104-S trimer complex. Representative micrograph, 2D class averages, gold-standard FSC plot, and local resolution estimation for **c-e,** C119-S2P and, **d-f**, C104-S. Both complexes revealed binding of Fabs to both “down” and “up” RBD conformations.

**Extended Data Figure 5.**
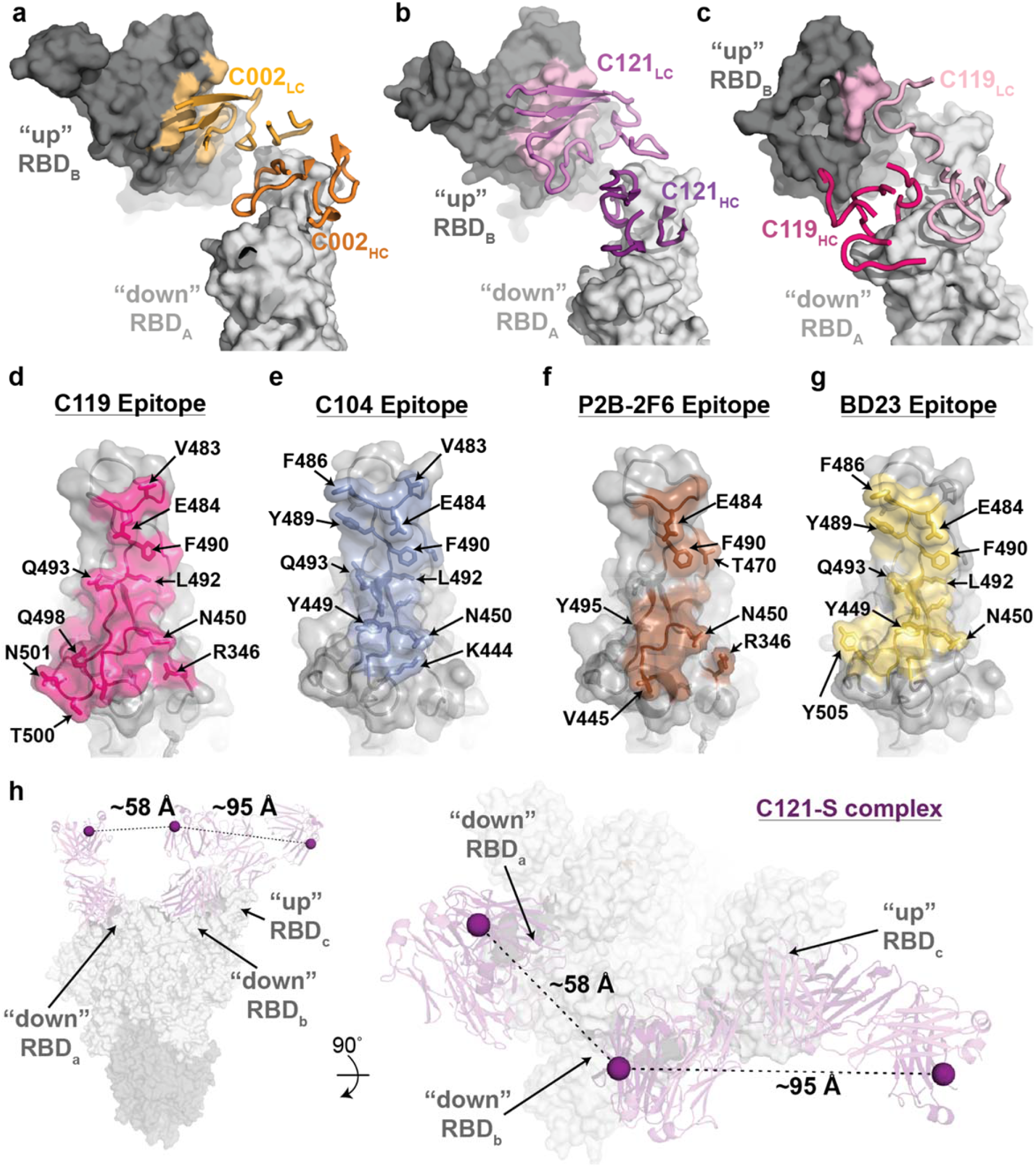
Primary and secondary epitopes of class 2 hNAbs. **a-c,** Primary epitopes for C002 (panel a), C121 (panel b), and C119 (panel c) on “down” RBD. A secondary epitope is observed if a Fab is bound to an adjacent “up” RBD for these NAbs. Antibody paratopes are represented as cartoons. A similar interaction in the C104-S structure is not shown due to low local resolution on the “up” RBD. **d-g**, Primary epitopes for C119 (panel d), C104 (panel e), P2B-2F6 (panel f; PDB 7BWJ), and BD23 (panel g, PDB 7BYR). The existence of secondary epitopes for P2B-2F6 and BD23 cannot be determined because the P2B-2F6 epitope was determined from a crystal structure with an RBD^6^, and the BD23-S cryo-EM structure showed only one bound Fab^8^. **h**, Measurement of Cα distance between the C-termini of adjacent C121 C_H_1 domains (residue 222_HC_ on each Fab). Measurements of this type were used to evaluate whether intra-spike crosslinking by an IgG binding to a single spike trimer was possible for hNAbs in Extended Data Table 1.

**Extended Data Figure 6.**
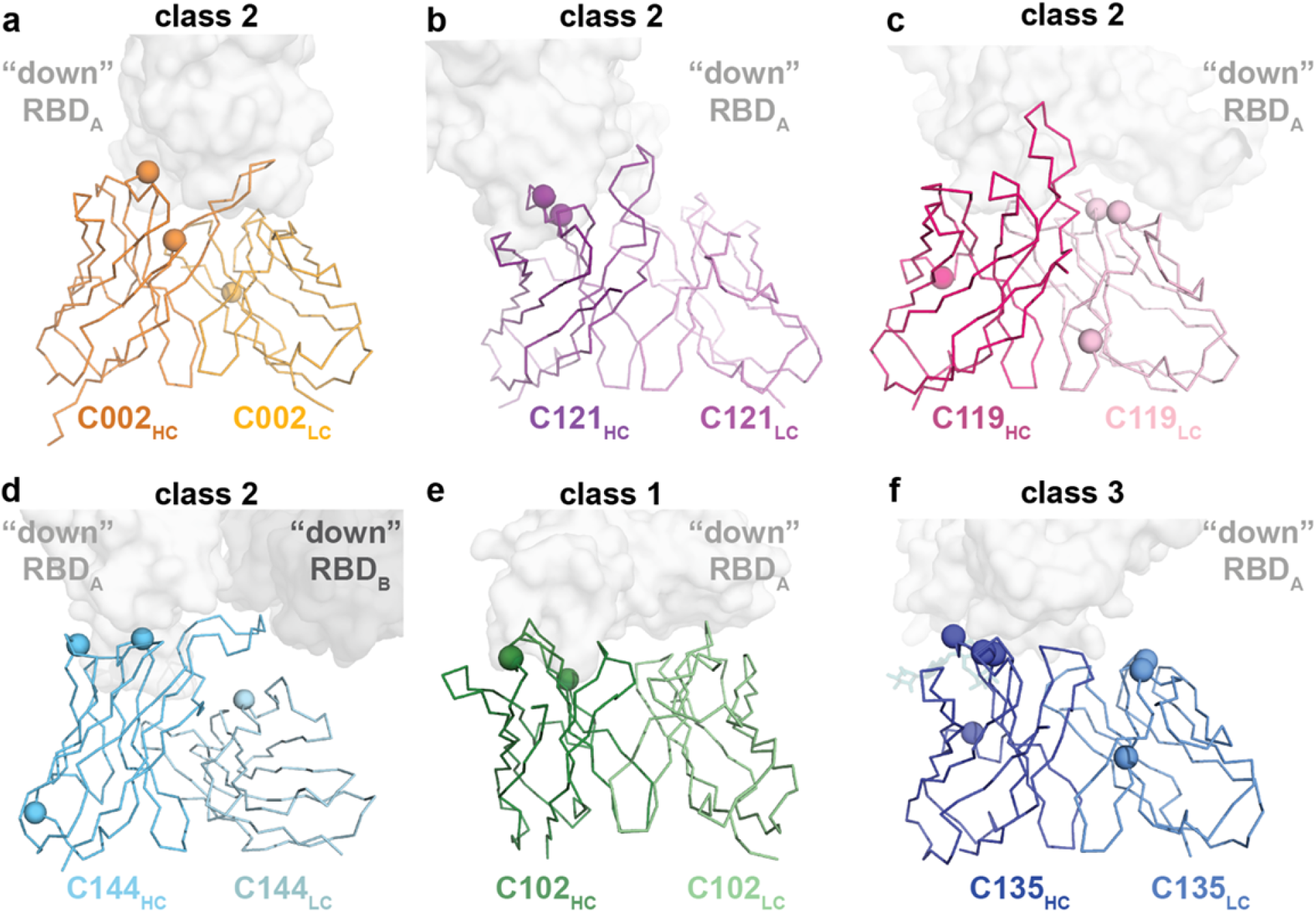
Mapping somatic hypermutations (SHMs) of SARS-CoV-2 NAbs. **a-f,** Somatic hypermutations in HC and LC V gene segments for C002 (panel a), C121 (panel b), C119 (panel c), C144 (panel d), C102 (panel e) and C135 (panel f) are shown as spheres on the antibody V_H_ and V_L_ domains (ribbon representations). The primary RBD epitope is shown as a light gray surface; secondary RBD epitope for C144 is in dark gray.

**Extended Data Figure 7.**
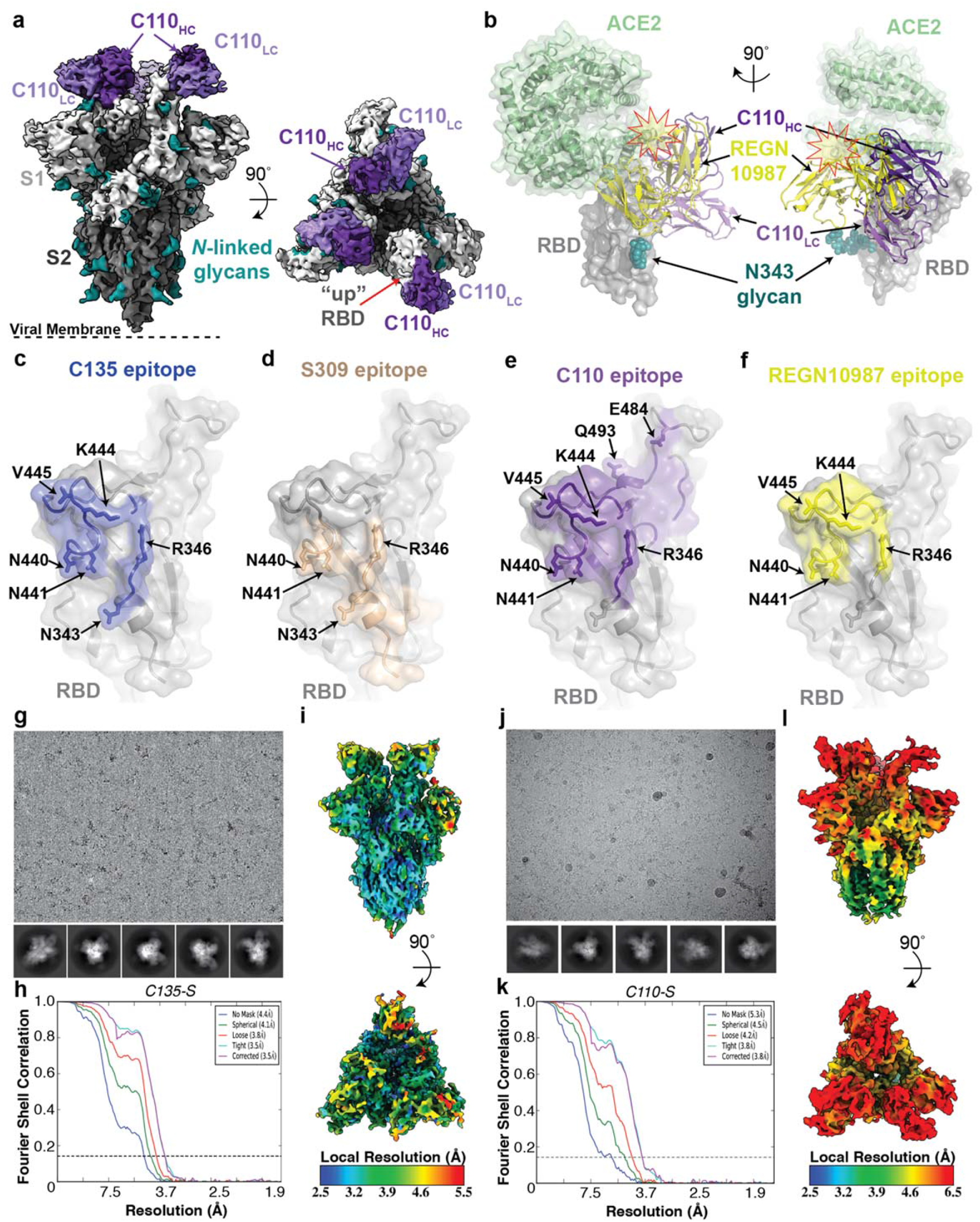
Cryo-EM structure of C110-S complex and epitope mapping. **a,** 3.8 Å cryo-EM reconstruction of C110-S trimer complex. **b,** Composite model of C110-RBD (purple and gray, respectively) overlaid with the SARS-CoV-2 NAb REGN-10987 (yellow, PDB 6XDG) and soluble ACE2 (green, PDB 6M0J). Model was generated by aligning structures on 188 RBD Cα atoms. **c-f,** Surface representation of RBD epitopes for **c,** C135 (blue), **d,** S309 (brown, PDB 6WSP), **e**, C110 (purple) and **f**, REGN-10987 (yellow, PDB 6XDG). Given the low resolution of the antibody-RBD interface, epitopes were assigned by selection of any RBD residue within 7 Å of any antibody Cα atom. Mutation sites found in sequence isolates^17^ (green) and in laboratory selection assays^18^ (red) are shown. Representative micrograph, 2D class averages, gold-standard FSC plot, and local resolution estimation for **g-i,** C135-S 2P and, **j-l**, C110-S 2P. Both complexes revealed binding of Fabs to both 2 “down”/1 “up” RBD conformations.

**Extended Data Figure 8.**
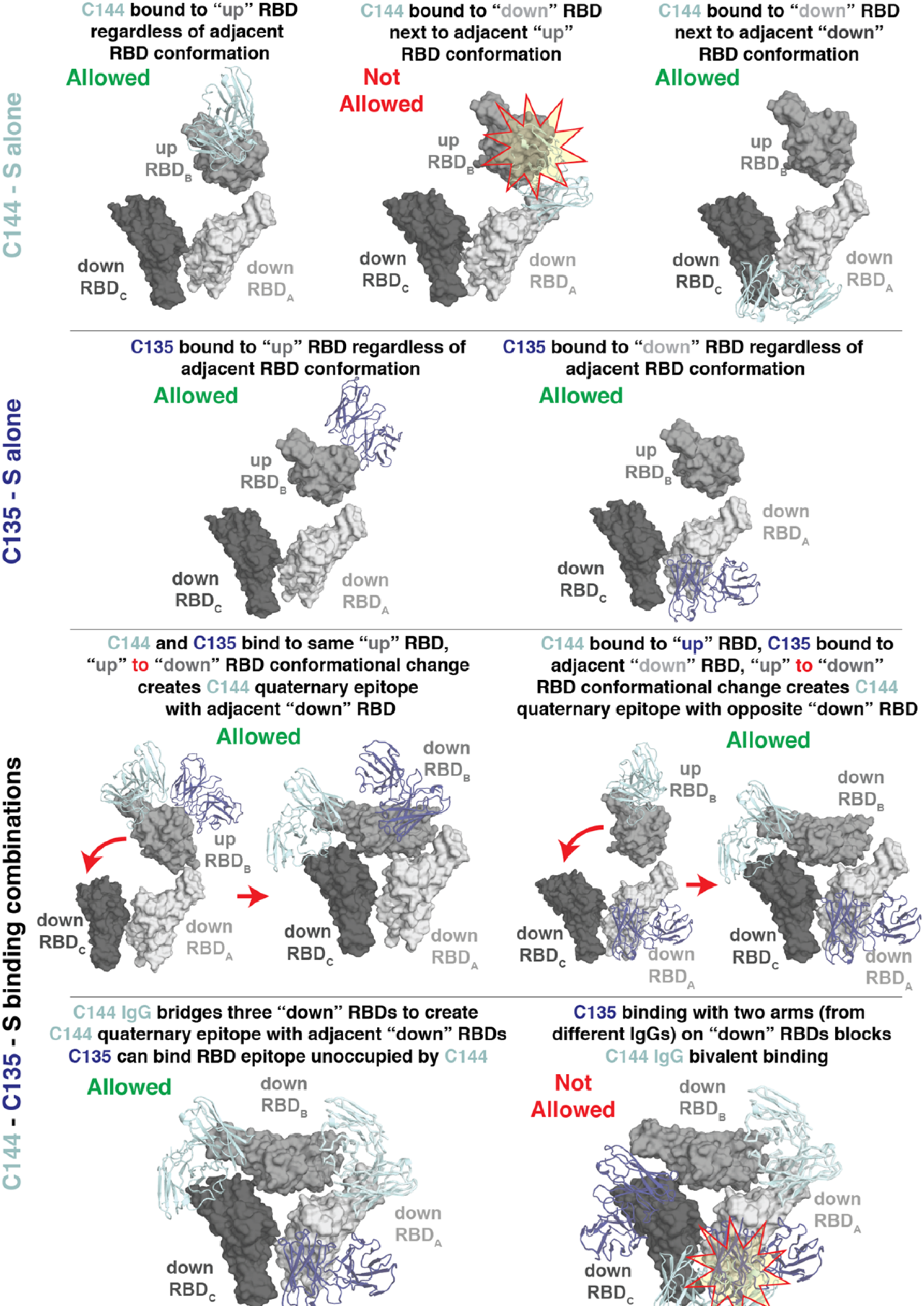
Possibilities for simultaneous engagement of C144 and C135 on spikes with different combinations of “up” and “down” RBDs. Modeling of C144 (light blue) and C135 (dark blue) V_H_-V_L_ domains on different RBD conformations. Steric clashes are shown as a red and yellow star.

**Extended Data Figure 9.**
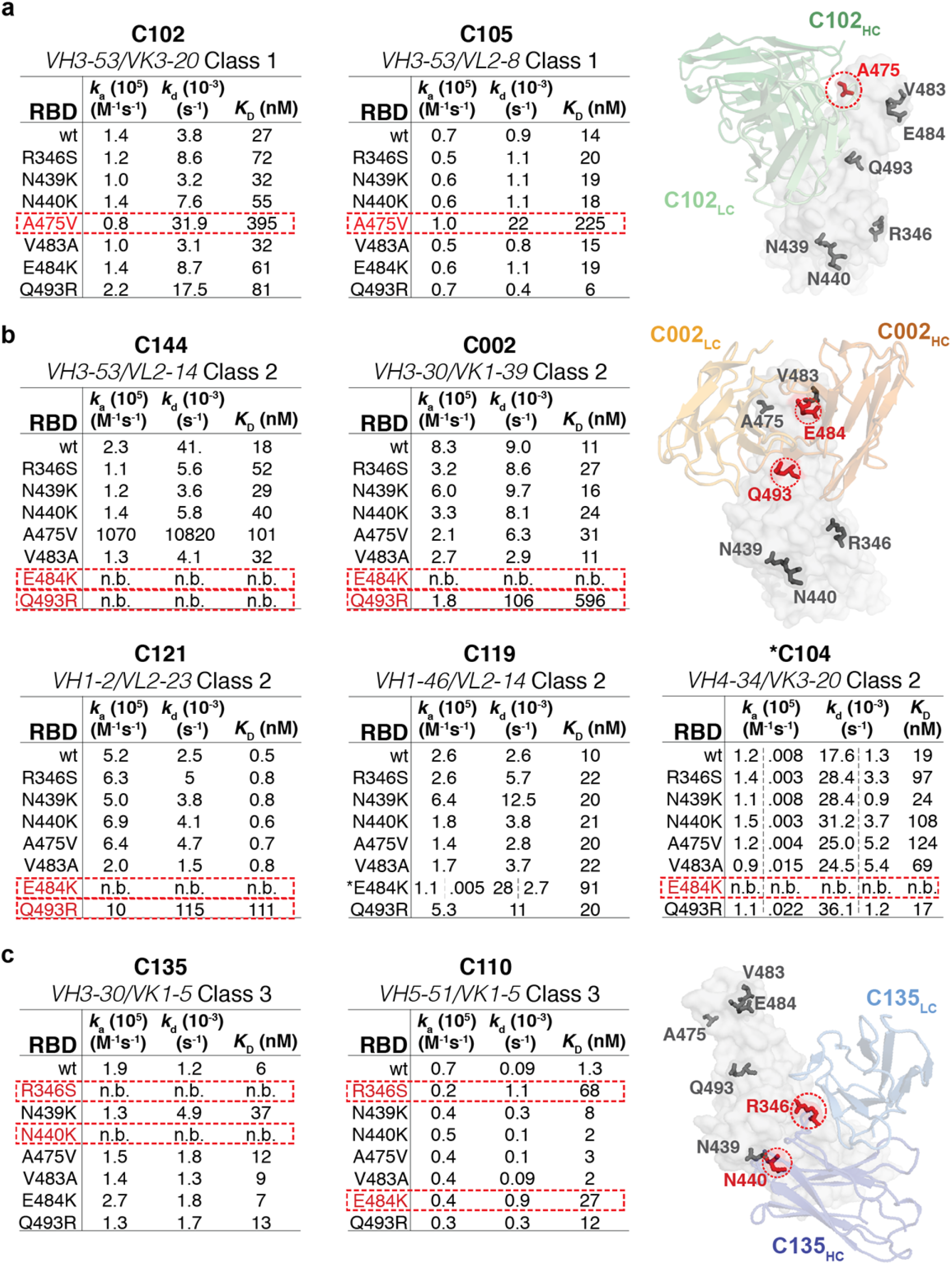
SPR binding data for hNAbs. Kinetic and equilibrium constants for binding to unaltered RBD (indicated as wt) and mutant RBDs are shown in tables beside structures of a representative NAb-RBD complex for each class. Residues that were mutated are highlighted as colored sidechains on a gray RBD surface. Antibody V_H_-V_L_ domains are shown as cartoons. Kinetic and equilibrium constants for NAbs that contact adjacent RBDs on S trimer (C144, C002, C119, and C121) do not account for contacts to a secondary RBD since binding was assayed by injected monomeric RBDs over immobilized IgGs. * indicates kinetic constants determined from a two-state binding model.

**Extended Data Figure 10:**
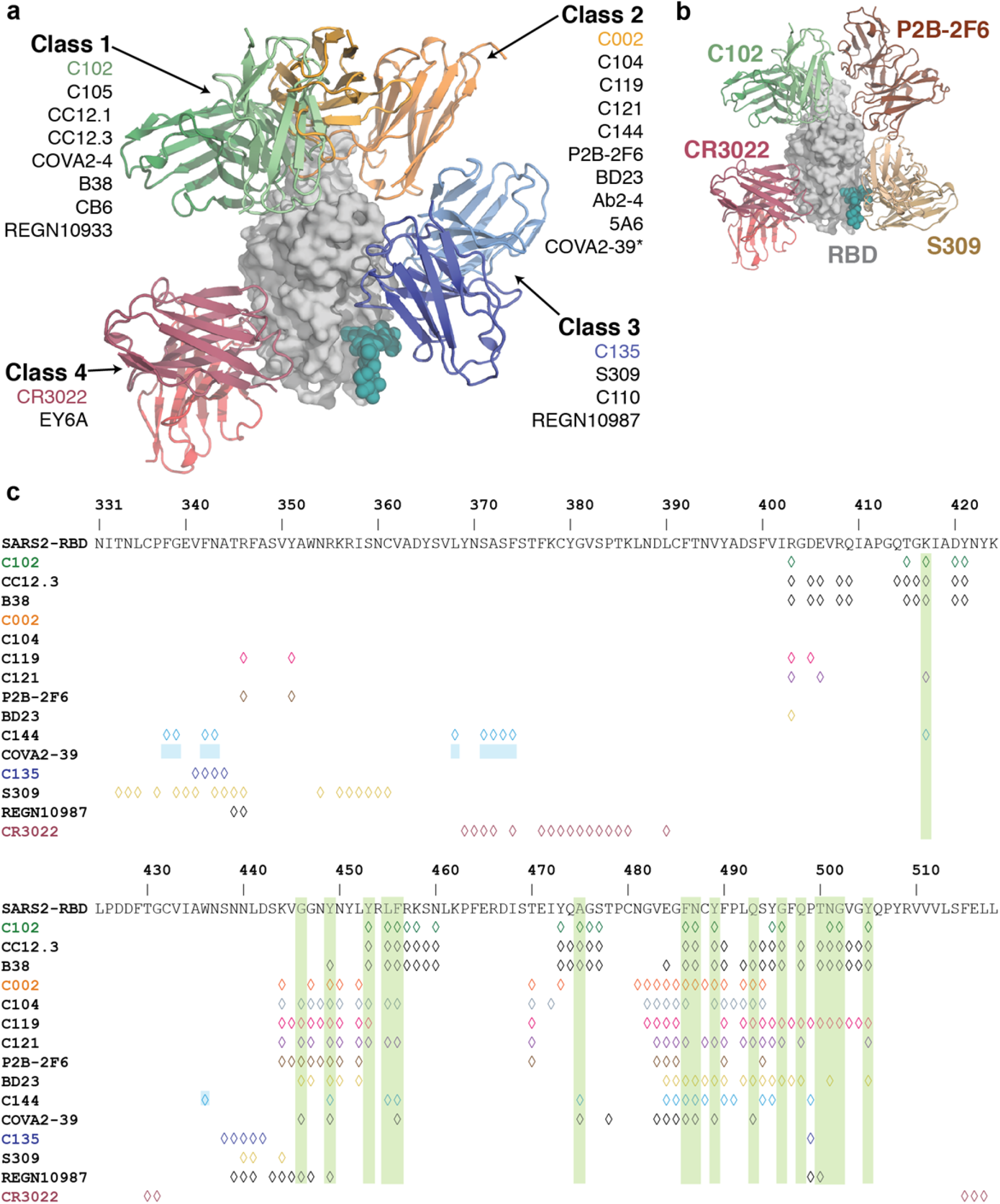
Summary of hNAbs. **a,** Structural depiction of a representative NAb from each class binding its RBD epitope. **b**, Composite model illustrating non-overlapping epitopes of NAbs from each class bound to a RBD monomer. **c**, Epitopes for SARS-CoV-2 NAbs. RBD residues involved in ACE2 binding are boxed in green. Diamonds represent RBD residues contacted by the indicated antibody.

**Extended Data Table 1.** Anti-SARS-CoV-2 NAb classification and structural properties.

| Antibody | Reference | IGHV<br>(# of aa<br>SHM) | CDRH3<br>length<br>(aa)<sup>†</sup> | IGLV<br>(# of aa<br>SHM) | CDRL3<br>length<br>(aa)<sup>†</sup> | IC<sub>50</sub>/IC<sub>90</sub><br>(ng/mL)<sup>†</sup> | Potential<br>IgG intra-spike<br>binding<sup>§</sup> | Contacts<br>adjacent<br>RBD | Structural Information |
| --- | --- | --- | --- | --- | --- | --- | --- | --- | --- |
| <u>Class 1: Blocks ACE2, accessibility of RBD epitope only in "up" conformation</u> |  |  |  |  |  |  |  |  |  |
| C102 | this study | VH3-53 (2) | 11 | VK3-20 (0) | 9 | 34 / 143 | ??? | ??? | 3.0 Å Fab-RBD |
| C105 | Barnes, et al.<sup>1</sup> | VH3-53 (0) | 12 | VL2-8 (1) | 11 | 26.1 / 134 | Yes | No | 3.4 Å Fab-S, PDB 6XCM |
| B38 | Wu, et al.<sup>2</sup> | VH3-53 (1) | 9 | VK1-9 (2) | 10 | 117 / NA | ??? | ??? | 1.8 Å Fab-RBD, PDB 7BZ5 |
| CC12.3 | Yuan, et al.<sup>3</sup> | VH3-53 (3) | 12 | VK3-20 (1) | 9 | 20 / NA | ??? | ??? | 2.9 Å Fab-RBD, PDB 6XC7 |
| <u>Class 2: Blocks ACE2, accessibility of RBD epitope in "up"/"down" conformations</u> |  |  |  |  |  |  |  |  |  |
| C002 | this study | VH3-30 (1) | 17 | VK1-39 (1) | 9 | 8.9 / 37.6 | Yes | Yes | 3.4 Å Fab-S |
| C104 | this study | VH4-34 (6) | 17 | VK3-20 (3) | 9 | 23.3 / 140 | Yes | Yes | 3.7 Å Fab-S |
| C119 | this study | VH1-46 (1) | 20 | VL2-14 (3) | 11 | 9.1 / 97.8 | Yes | Yes | 3.5 Å Fab-S |
| C121 | this study | VH1-2 (2) | 22 | VL2-23 (0) | 10 | 6.7 / 22.3 | Yes | Yes | 3.6 Å Fab-S |
| C144 | this study | VH3-53 (3) | 25 | VL2-14 (1) | 10 | 6.9 / 29.7 | Yes | Yes | 3.3 Å Fab-S |
| COVA2-39 | Wu, et al.<sup>4</sup> | VH3-53 (3) | 17 | VL2-23 (1) | 10 | 36 / NA | ??? | ??? | 1.7 Å Fab-RBD, PDB 7JMP |
| 5A6 | Wang, et al.<sup>5</sup> |  |  |  |  | 75.5 / NA | Yes | Yes | 2.4 Å Fab-S |
| P2B-2F6 | Ju, et al.<sup>6</sup> | VH4-38*02 (2) | 20 | VL2-8 (0) | 10 | 50 / NA | ??? | ??? | 2.9 Å Fab-RBD, PDB 7BWJ |
| Ab2-4 | Liu, et al.<sup>7</sup> | VH1-2 (3) | 15 | VL2-8 (0) | 10 | 394 / NA | Yes | No | 3.2 Å Fab-S, PDB 6XEY |
| BD23 | Cao, et al.<sup>8</sup> | VH7-4*02 (0) | 19 | VK1-5*03 (0) | 9 | 4800 / NA | No | No | 3.8 Å Fab-S, PDB 7BYR |
| <u>Class 3: Does not overlap with ACE2 binding site, accessibility of RBD epitope in "up"/"down" conformations</u> |  |  |  |  |  |  |  |  |  |
| C135 | this study | VH3-30 (4) | 12 | VK1-5 (3) | 9 | 16.6 / 48.9 | No | No | 3.5 Å Fab-S |
| S309 | Pinto, et al.<sup>9</sup> | VH1-18 (6) | 20 | VK3-20 (3) | 8 | 79* / NA | No | No | 3.1 Å Fab-S, PDB 6WPS |
| C110 | this study | VH5-51 (2) | 21 | VK1-5 (3) | 9 | 18.4 / 77.3 | No | No | 3.8 Å Fab-S |
| REGN10987 | Hansen, et al.<sup>10</sup> | VH3-30 (4) | 13 | VL2-14 (6) | 10 | 6.1 / NA | ??? | ??? | 3.9 Å Fab-RBD, PDB 6XDG |
| <u>Class 4: Does not overlap with ACE2 binding site, accessibility of RBD epitope only in "up" conformation</u> |  |  |  |  |  |  |  |  |  |
| CR3022 | Yuan, et al.<sup>11</sup> | VH5-51 (8) | 12 | VK4-1 (3) | 9 | >10,000 / NA | ??? | ??? | 3.1 Å Fab-RBD, PDB 6W41 |
| COV1-16 | Liu, et al.<sup>12</sup> | VH1-46 (1) | 20 | VK1-33 (3) | 10 | 130 / NA | ??? | ??? | 2.9 Å Fab-RBD |
| EY6A | Zhou, et al.<sup>13</sup> | VH3-30*18 (3) | 14 | VK1-39 (0) | 10 | 70-20,000** / NA | No | Yes | 3.7 Å Fab-S, PDB 6ZDH |

<sup>†</sup>Average human antibody CDRH3 and CDRL3 lengths are 15 (CDRH3) and 9-10 (CDRL3) amino acids.
\*IC<sub>50</sub> calculated against authentic SARS-CoV-2 virus.
\*\*IC<sub>50</sub> varied depending on neutralization assay utilized.
<sup>†</sup>Unknown IC<sub>90</sub>s indicated as NA (not available).
<sup>§</sup>Potential for intra-spike crosslinking by an IgG binding to a single spike trimer was evaluated as described in the Methods.
?? Inference that cannot be made from a structure of a Fab bound to a RBD.
IGHV = Immunoglobulin heavy chain variable gene segment;
IGLV = Immunoglobulin light chain variable gene segment
V gene segments, somatic hypermutation (SHM) information, CDR lengths, IC<sub>50</sub>/IC<sub>90</sub> values for NAbS in this study are from ref.<sup>14</sup>.

